# An absolute quantification atlas of small non-coding RNAs across diverse mammalian tissues and cell lines

**DOI:** 10.1101/2025.07.31.667863

**Authors:** Wen Xiao, Yuli Zheng, Hongdao Zhang, Beiying Xu, Ruiwen Zhang, Ligang Wu

**Affiliations:** Key Laboratory of RNA Science and Engineering, Shanghai Key Laboratory of Molecular Andrology, Shanghai Institute of Biochemistry and Cell Biology, Center for Excellence in Molecular Cell Science, Chinese Academy of Sciences, University of Chinese Academy of Sciences, Shanghai, 200031, China

## Abstract

The low quantitative accuracy of conventional small noncoding RNA sequencing (sncRNA-seq) methods due to extensive ligation bias commonly limits functional investigation of microRNAs (miRNAs) and PIWI-interacting RNAs (piRNAs). Here, we develop 4NBoost, a single-tube sncRNA-seq protocol designed to minimize bias in the estimated absolute quantification of miRNA and piRNA transcripts through the incorporation of quantitative exogenous RNA spike-ins. With 4NBoost, we profile sncRNA expression across 20 murine tissues, 18 macaque tissues, and 24 widely used cell lines, as well as 4 *Arabidopsis* tissues, to establish a comprehensive quantitative reference atlas. Compared with existing small RNA databases, our data reveal substantial biases in miRNA abundance, strand selection, and tissue-specific expression at both individual and family levels. To further extend its utility, we employ machine learning to model and correct biases in conventional datasets, effectively recovering ground truth transcript abundances. All 4NBoost data and the accompanying bias-correction model are freely available via SmRNAQuant (http://wulg-lab.sibcb.ac.cn/SmRNAQuant/), a web-based repository for exploring sncRNA expression. Together, the 4NBoost, bias-correction model, and SmRNAQuant provide powerful resources to advance sncRNA research.

## INTRODUCTION

Small non-coding RNAs (sncRNAs), including microRNAs (miRNAs), small interfering RNAs (siRNAs), and piwi-interacting RNAs (piRNAs), are key regulators in a variety of biological and pathological processes such as organ development, epigenetic modification, and tumorigenesis [1–8]. Numerous recent studies have greatly expanded our understanding of the diversity, complexity, and function of these sncRNAs [9–19]. Given their critical regulatory roles, a robust and comprehensive understanding of their biological functions requires accuracy in sncRNA profiling and quantification [20]. Despite the power and versatility of next-generation sequencing tools for studying sncRNA expression [21, 22], conventional sncRNA library construction typically relies on T4 RNA ligase for adapter ligation prior to reverse transcription and amplification, which are prone to introducing biases in quantification [23–26]. These biases primarily arise from the structure-dependent mechanism of T4-mediated ligation of adapters to sncRNAs, causing certain sncRNAs to be over- or under-represented, which can ultimately skew conclusions about their biological functions [27–31].

Several methods have been developed to reduce or minimize these biases, including the use of terminal randomized adapter sequences and/or high polyethylene glycol (PEG) concentrations, as reported in methods such as 4N-seq, AQ-seq, IsoSeek, and NEXTflex [9, 15, 29–35]. Despite these advances, the large majority of sequencing approaches primarily target miRNAs with a 2’-hydroxyl group (2’-OH) at the 3’ end, which neglects optimization of ligation efficiency for piRNAs and plant miRNAs, which instead harbor 3’ 2’-O-methyl modifications (2’-Ome) that hinder ligation [15, 36–39]. Furthermore, many of these methods do not incorporate unique molecular identifiers (UMIs) to correct for PCR bias by eliminating duplicate sequences. Additionally, few of these sequencing approaches has been adopted to determine estimated absolute quantification of sncRNAs, which is essential for addressing crucial fundamental research questions related to sncRNA stoichiometry. For example, miRNA activity has been shown to depend on a threshold level of expression, with only the most abundant miRNAs capable of effectively mediating target suppression [40–44]. This indicates that intracellular concentration is tightly coupled to functional activity. However, absolute expression levels have been measured for only a limited subset of miRNAs in specific cell or tissue types [43–47]. Additionally, understanding the precise miRNA concentrations required for regulatory function can be vital to identifying differences between physiological and pathological processes [48, 49]. Moreover, existing miRNA databases—such as miRbase [50], MirGeneDB [51], miRmine [52], and DIANA-miTED [53]—as well as tissue atlas of sncRNAs [54, 55], were constructed using such biased sncRNA sequencing data, and therefore lack the capacity for accurate absolute quantification. Consequently, a comprehensive, bias-minimized, and quantitatively accurate reference for sncRNAs has yet to be established.

To overcome these challenges, in the current study, we developed 4NBoost, a quantitative miRNA and piRNA sequencing technique that minimizes bias in small RNA sequencing through the incorporation of exogenous ratiometric RNA spike-ins. We applied 4NBoost to generate a comprehensive reference atlas of sncRNA expression by profiling 259 samples, including 20 tissues from mice, 18 tissues from crab-eating macaques, 24 commonly used cell lines, and 4 tissues from *Arabidopsis*. To the best of our knowledge, this is the most systematic estimated absolute quantification of sncRNAs across mammalian tissues and cell lines. To further expand its utility, we developed an XGBoost-based framework to correct biases in conventional datasets, enabling accurate reconstruction of ground truth transcript abundances. Additionally, we developed a web-based database to facilitate access to 4NBoost data, allowing researchers to retrieve and analyze small RNA expression profiles. Together, the 4NBoost method, correction model, and online resource constitute a robust platform for advancing both fundamental and clinical research in small noncoding RNAs.

## RESULTS

### Construction of 4NBoost with greatly reduced ligation bias for high accuracy sncRNA quantification

To construct 4NBoost, we modified the 4N-Xu method [35, 56] by first increasing the size of random nucleotides (NTs) in 5’ adapter from 4 to 7 NTs, while retaining a 3’ adapter of 4 random NTs that ensures a total of 11 random bases to reduce ligation bias and expand the number of unique molecular identifiers (UMIs) essential for removing duplicate reads. Based on AmpUMI model predictions [57] and previous studies [34], we determined that UMIs with 11 random bases could effectively capture the most abundant molecules with a low probability of UMI collision (1.4%, Supplementary Figure 1A). To limit the likelihood of generating excessive byproducts associated with long consecutive random sequences, we also introduced 3 fixed nucleotides into the 7 random nucleotides of the 5’ adapter [58].

We then designed and synthesized 30 spike-ins, including 7 with 2’-O-methyl modifications at the 3’ end, which enabled the evaluation of the quantitative accuracy of 2’-O-methyl-modified sncRNAs in addition to miRNAs. We prepared two spike-in pools, including an equimolar pool (EM) and a ratiometric pool (RM), to test the accuracy and uniformity of the library (Supplementary Figure 1B). In the ratiometric pool, the concentrations of 30 spike-ins spanned an approximate 2 x 10^5^-fold concentration range, with 3 oligos per concentration.

Optimization of reaction conditions showed that, in addition to high PEG8000 concentration [32], the coefficient of variation (CV) of equimolar spike-ins gradually decreased with increasing concentration of adapters, suggesting that the higher concentration of adapters could further minimize potential library biases (Supplementary Figure 1C). However, higher adapter concentrations led to a significant increase in self-ligation byproducts, with up to 90% of raw data comprising such artifactual sequences that had to be discarded (Supplementary Figure 1D). To balance library uniformity and data quality, we determined that 0.55 µM was the optimal concentration of 3’ and 5’ adapters (Supplementary Figure 1E). Additionally, we found that Lambda exonuclease was more efficient than RecJ exonuclease in depleting excess 3’ adapters [59, 60] (Supplementary Figure 1F-G).

We then evaluated the accuracy of 4NBoost by adding the EM or RM pools to the total cellular RNAs and calculating the relative amounts of spike-ins based on the sequencing results. Compared to TruSeq and AQ-seq, 4NBoost demonstrated the lowest average CV (1.73, 0.56, and 0.51, respectively, for TruSeq, AQ-seq, and 4NBoost), indicating a marked reduction in library ligation biases and resulting in more uniform representation of equimolar spike-ins (Figure 1A-B and Supplementary Figure 1H). Correlation analysis between the expected and observed abundance of ratiometric spike-ins yielded Pearson’s coefficients > 0.94, indicating that 4NBoost could accurately reflect small RNA expression levels across varying abundances (Figure 1C). Notably, in 4NBoost libraries, spike-ins with 3’ end methylation showed similar ligation efficiency to non-methylated spike-ins (Supplementary Figure 1I), in sharp contrast to the TruSeq libraries, where spike-ins with terminal methylation were detected at over 10-fold lower efficiency compared to their non-methylated counterparts. This finding was further validated by the 4NBoost library from mouse testis, which displayed a stronger 1U signal for piRNAs compared to the TruSeq library (Supplementary Figure 1J).

**Figure 1:**
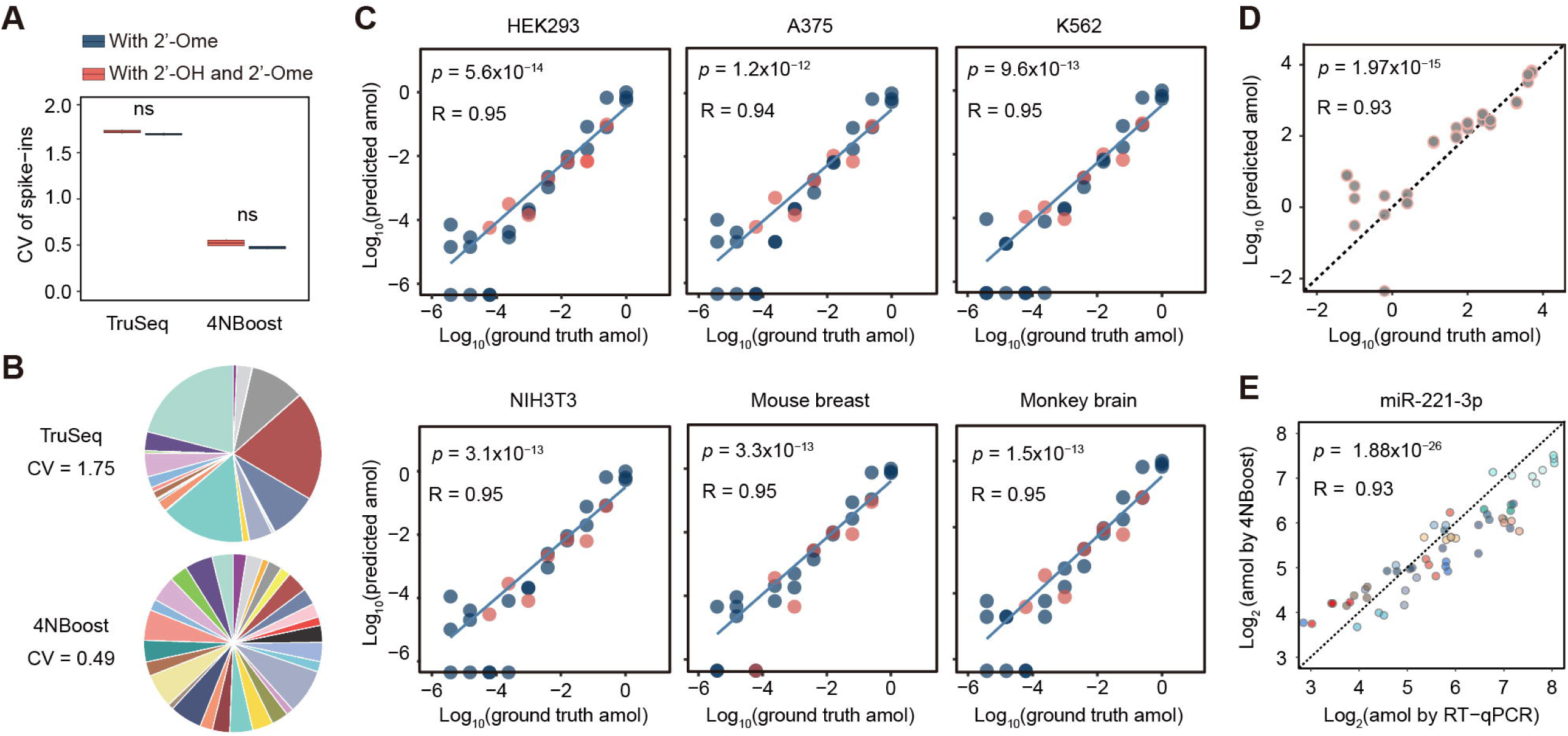
4NBoost significantly reduces ligation bias and achieves accurate quantification. **(A)** The CV of the distribution of 30 (sequences with 2’-OH or 2’-Ome at the 3’ ends) or 7 (sequences with 2’-Ome at the 3’ ends) equimolar spike-ins, using total HEK293 cellular RNA as background, is shown in the box plots. In these plots, the center line represents the median, while the box edges correspond to the 25th and 75th percentiles. The whiskers extend to the minimum and maximum values. The biological replicates for these experiments were n = 2 for TruSeq and n = 4 for 4NBoost. **(B)** Pie charts illustrating the proportion of 30 equimolar spike-ins detected by TruSeq and 4NBoost in representative samples. **(C)** The correlations between the ground truth and predicted abundances of ratiometric spike-ins in various samples are illustrated, with blue points representing unmodified sequences and red points indicating sequences with 2’-Ome at the 3’ ends. **(D)** Scatter plots illustrating the correlation between the ground truth and predicted abundances of additional external spike-ins. **(E)** Scatter plot illustrating the correlation between the predicted abundances of the selected miRNA as measured by 4NBoost and RT-qPCR, with different colors representing distinct cell lines. The number of biological replicates per cell line was three for Caco2 and four for all other cell lines. All p-values were calculated using two-tailed Student’s t-tests. Source data are provided as a Source Data file.

To assess the sensitivity of 4NBoost, we performed sequencing on serial dilutions of input RNA, ranging from microgram (µg) to nanogram (ng) and picogram (pg) amounts. As a benchmark, miRNA expression profiles generated from 1 µg of input RNA were compared with those from lower input amounts (Supplementary Figure 1K–L). The number of detected miRNA species remained relatively constant down to 10 ng of input RNA. At 1 ng, reproducibility in miRNA detection declined moderately, but Spearman correlation coefficients remained high (> 0.7). Below 1 ng, both the number of detected miRNA species and robustness decreased sharply. These results indicate that 4NBoost can faithfully profile sncRNA expression from as little as 1 ng of total RNA without substantial loss of performance.

Finally, we applied 4NBoost to quantify each small RNA by generating standard curves using the ratiometric spike-ins with known absolute molecule numbers. To validate its accuracy, we incorporated four additional external spike-ins and found that 4NBoost could predict the molar quantities of all spike-ins, except the lowest abundance one, with high precision (Pearson’s coefficient = 0.93; predicted vs ground truth values; Figure 1D). Additionally, a comparison between miR-221-3p abundance in ten cell lines predicted by 4NBoost and its copy numbers obtained by RT-qPCR showed a high correlation (Pearson’s coefficient = 0.93) between methods (Figure 1E). Similarly, miR-21-5p, the most abundantly expressed miRNA across most cell lines, showed excellent agreement between RT-qPCR and 4NBoost measurements (Pearson’s coefficient = 0.99, p = 4.95×10^-20^; Supplementary Figure 1M). Collectively, these results further support the high quantitative accuracy of our method. Notably, 4NBoost could accurately quantify sncRNAs with abundances >10 amol per 100 ng total RNAs.

### Construction of a quantitative sncRNA expression atlas of mammalian tissues and cell lines

Assembling a set of tissues and cell lines commonly used in miRNA studies (Supplementary Data 1) resulted in a panel of 24 cell lines (n = 4), 20 tissues from BALB/C mice (n = 4), and 18 tissues from crab-eating monkeys (n = 4) (Figure 2A). Using 4NBoost, we then constructed 244 total libraries, among which 242 passed quality control screening (Figure 2B). Each library contained over 1 million genome-mapped reads (Supplementary Figure 2A), which were further annotated into various small ncRNA species (Figure 2B). Among these ncRNAs, miRNA was the most well-represented species in most cell lines and tissues, except in testis samples (Figure 2C). Correlation analysis of miRNAs between biological replicates showed expression correlations exceeding 0.9 (Supplementary Figure 2B-D), indicating high reproducibility. The high correlation of the read counts before and after UMI deduplication for the top 100 most abundant miRNAs further supports that 11-nt UMIs are sufficient for accurate deduplication with negligible impact from potential UMI collisions (Supplementary Figure 2E).

**Figure 2:**
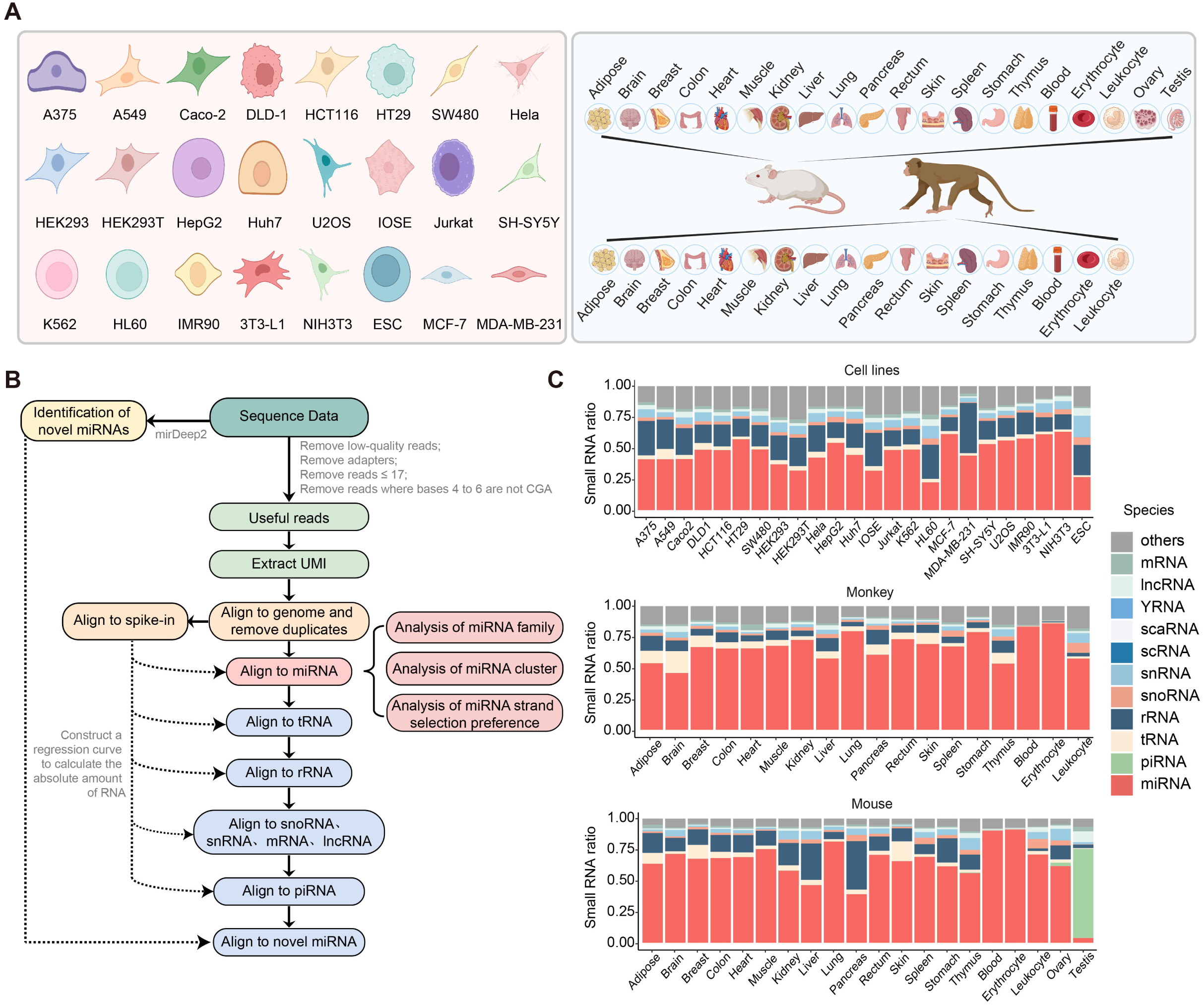
Overview of miRNA expression across various tissues and cell lines. **(A)** Tissues and cell lines collected for this study. This schematic was created with BioRender. **(B)** Data analysis workflow for this study. **(C)** Proportions of various small RNA types within all small RNA libraries. Biological replicates: n = 2 (for mouse ovary and testis), n = 3 (for Caco2, mouse-breast, monkey-rectum, and monkey-stomach), or n = 4 (for remaining samples).

Subsequent validation of quantitative accuracy for sncRNAs in each sample with an additional four spike-ins yielded a correlation coefficient of 0.97 between predicted and ground truth values (Supplementary Figure 2F-G), thereby confirming the accuracy of 4NBoost. This high accuracy enabled further estimation of the abundance of each miRNA and piRNA in all cell lines and tissues in our panel, which we used to construct a dataset of more reliable expression levels. We also noted that miRNA family quantifications had markedly low CV across biological replicates, further supporting the stability and reliability of quantification by 4NBoost (Supplementary Figure 2H). Overall, these results indicated that 4NBoost generated robust, quantitative miRNA and piRNA expression datasets across a wide range of tissues and cell lines.

### Comparative analysis of miRNA species and expression across cell lines and tissues

We then applied these quantitative sncRNA datasets to analyze the number of miRNA species expressed in each cell line and tissue, mitigating the influence of variations in sequencing depth by randomly selecting 1.5 million mapped molecules from each sample for analysis. In most cell lines, approximately 600 miRNA species were detected (Supplementary Data 2, Figure 3A), which was comparable to the 532 average miRNA species reported in the miRmine database [52]. Notably, U2OS cells had the highest number of miRNAs (715 species), while the fewest were detected in MDA-MB-231 cells (266 species). In tissue samples, typically more than 500 miRNA species were expressed, with the exception of erythrocytes and blood, which had only 328 species (Figure 3B). Of note, some samples with lower miRNA species numbers, e.g., erythrocytes and blood, also had a small subset of predominant or overrepresented miRNAs, such as miR-451. Intriguingly, we also observed that most samples from monkeys had fewer miRNA species compared to the corresponding tissues from mice, likely due to the less comprehensive miRNA annotation in monkeys. However, detected miRNA numbers may be influenced by factors such as sncRNA composition or incomplete annotation, and should be interpreted with caution.

**Figure 3:**
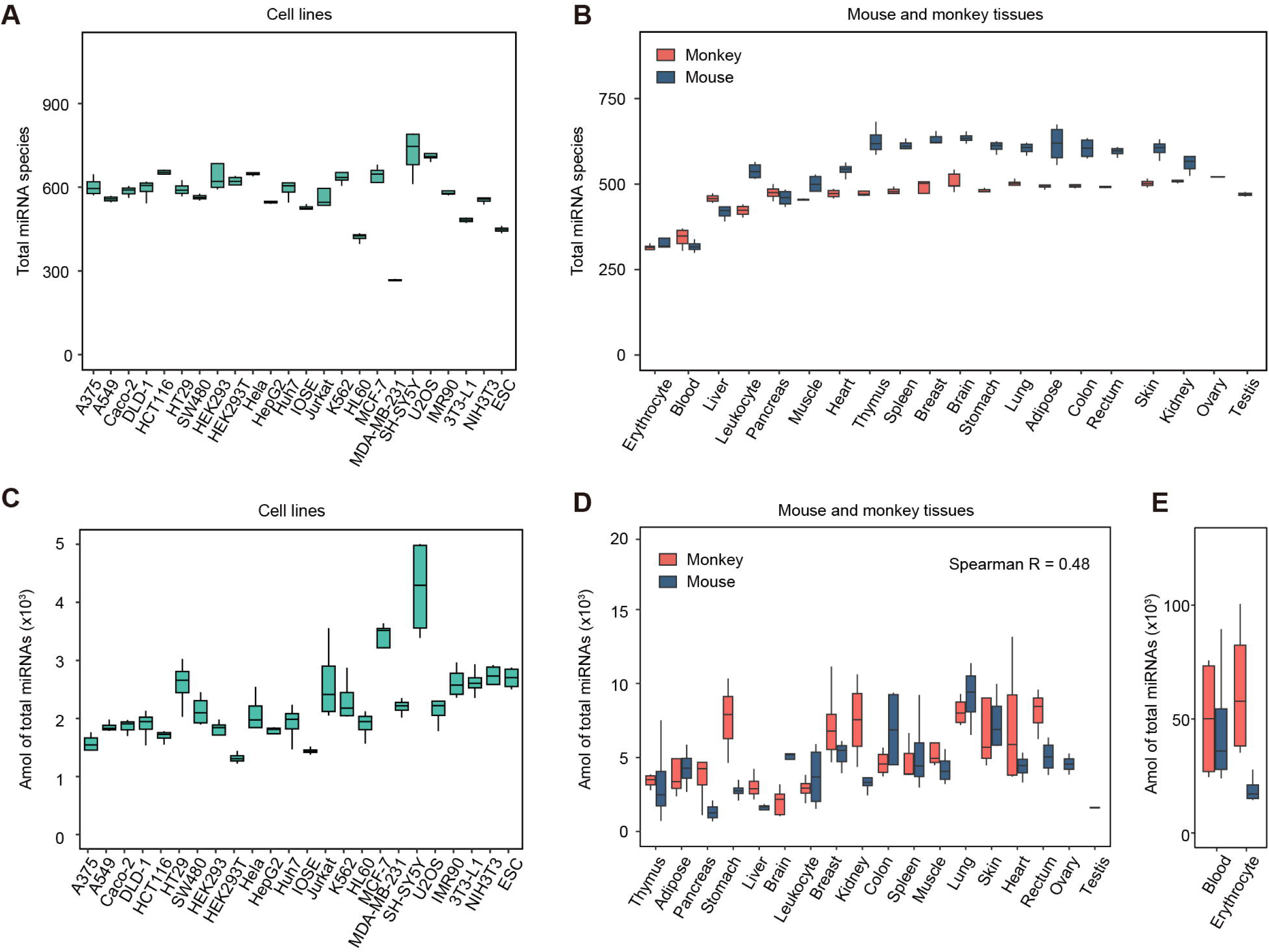
Species and quantification of miRNAs across cell lines and tissues. (A-B) Box plots showing the number of detected miRNA species across various cell lines **(A)** and tissues **(B)**, with 1.5 million mapped molecules from each sample used for analysis. **(C-E)** Box plots illustrating the quantification of miRNA in various cell lines **(C)** and tissues **(D-E)**. In these plots, the center line represents the median value, the box borders represent the upper and lower quartiles (25th and 75th percentiles, respectively), and the ends of the top and bottom whiskers represent maximum and minimum values, respectively. Biological replicates for (A-E): n = 2 (for mouse ovary and testis), n = 3 (for Caco2, mouse-breast, monkey-rectum, and monkey-stomach), or n = 4 (for the remaining samples).

We then compared miRNA concentrations quantified by 4NBoost (Figure 3C-E). Similar to the distribution of miRNA species across samples, total miRNA contents were comparable among most cells, with an average of ∼2200 amol per 100 ng of total RNAs (Supplementary Data 2, Figure 3C), or approximately 15 pg (∼0.015% of total RNAs), which was consistent with previous reports [61]. Across cell lines, SH-SY5Y cells had the highest miRNA expression level, ∼4000 amol per 100 ng total RNA. Across tissues, for example, in monkeys, we found that brain samples had the lowest average miRNA levels, while heart and lung samples had the highest. By contrast, in mice, pancreas and testis tissues had the lowest average miRNA levels, whereas lung and skin samples had the highest miRNA contents (Supplementary Data 2, Figure 3D). As a result of this variability, Spearman analysis of miRNA concentrations between monkey and mouse tissues showed low correlation (R = 0.48; Figure 3D), potentially reflecting big differences in miRNA expression among species. Additionally, due to the overrepresentation of miR-451 in blood and erythrocyte samples, miRNA quantities were much higher in these samples compared to solid tissues (Figure 3E).

Notably, although pancreas samples had a higher relative proportion of miRNAs than testis in mice (Figure 2C), their quantities were not significantly different (Figure 3D). These findings highlighted the importance of absolute quantification for comparing miRNA expression levels across different tissues or cell lines.

### Improved miRNA expression accuracy with 4NBoost

Previous studies have shown that miRNA dosage influences the strength of downstream gene regulation, with highly expressed miRNAs commonly selected as key targets for further investigation [48, 49]. In our study, we focused on these highly expressed miRNAs across various cell lines and tissues, identifying notable discrepancies between 4NBoost datasets and the DIANA-miTED database, which shares the greatest overlap in cell lines with our study, as well as MirGeneDB, a well-established source for miRNA expression data in tissues. Estimated absolute quantification with 4NBoost suggested that the expression levels of numerous miRNA species, including pre-mir-19, pre-mir-29, pre-mir-23a, pre-mir-24, pre-mir-126a, pre-mir-143, pre-mir-26a, and pre-mir-30c were likely underestimated in DIANA-miTED and MirGeneDB, while pre-let-7a, pre-let-7b, pre-let-7c, pre-mir-10a, pre-mir-22, and pre-mir-191 were overestimated in samples from the same tissues or cell lines (Supplementary Data 3).

To test whether these discrepancies arose through differences among library construction methods, we analyzed heterodimer structures for both over- and underestimated miRNAs, as the influence of heterodimer structures formed between RNA and adapters has been widely reported to influence T4 RNA ligase (Rnl) activity, wherein 5’ adapter ligation by Rnl1 favoring RNAs with an unpaired 5’ end [29, 30, 62–64]. This analysis of heterodimer structures in over- and underestimated miRNAs with 5’ RNA adapters revealed that over 85% of the overestimated miRNAs had unpaired 5’ ends, while around 70% of underestimated miRNAs had paired 5’ ends (Supplementary Figure 3A-B), which was consistent with Rnl1 bias for unpaired 5’ ends. By contrast, for 4NBoost data, generated with random adapters, over 90% of these underestimated miRNAs formed secondary structures more favorable for Rnl1 ligation (Supplementary Figure 3C). These findings suggested that 4NBoost consequently improved the accuracy of miRNA expression quantitation, compared to that in DIANA-miTED or MirGeneDB.

### Analysis of miRNA strand preference by 4NBoost

In addition to the above problem of over- or underrepresentation, ligation bias has also been identified as a potential source of mis-annotation in miRNA strand preference for certain miRNAs, such as pre-mir-423, pre-mir-17, pre-mir-106b, and pre-mir-151a. As bias-reducing methods have been shown to help mitigate this issue [29, 32], we next examined strand preference in 4NBoost data. This analysis showed that miRNA strand preferences were consistent with previously validated results for pre-mir-324 [65], pre-mir-423 [32], pre-mir-223 [66], and pre-mir-133a [67] (Figure 4A, Supplementary Table 1). Moreover, strand preferences measured by 4NBoost also correlated closely with those obtained by AQ-seq in the HCT116 and HEK293T cell lines (Pearson’s coefficient = 0.94 and 0.91, respectively), whereas moderate differences were noted when compared to TruSeq (Pearson’s coefficient = 0.79 for the HCT116 cell line and 0.82 for the HEK293T cell line; Figure 4B). Subsequent application of a strand selection model to all miRNA, as previously described [68], and compared with the strand ratio data obtained by 4NBoost and TruSeq showed that 4NBoost data shared a stronger correlation with model predictions (Pearson’s coefficient = 0.67) than TruSeq (Pearson’s coefficient = 0.59; Figure 4C), further emphasizing higher accuracy of 4NBoost in strand ratio measurements.

**Figure 4:**
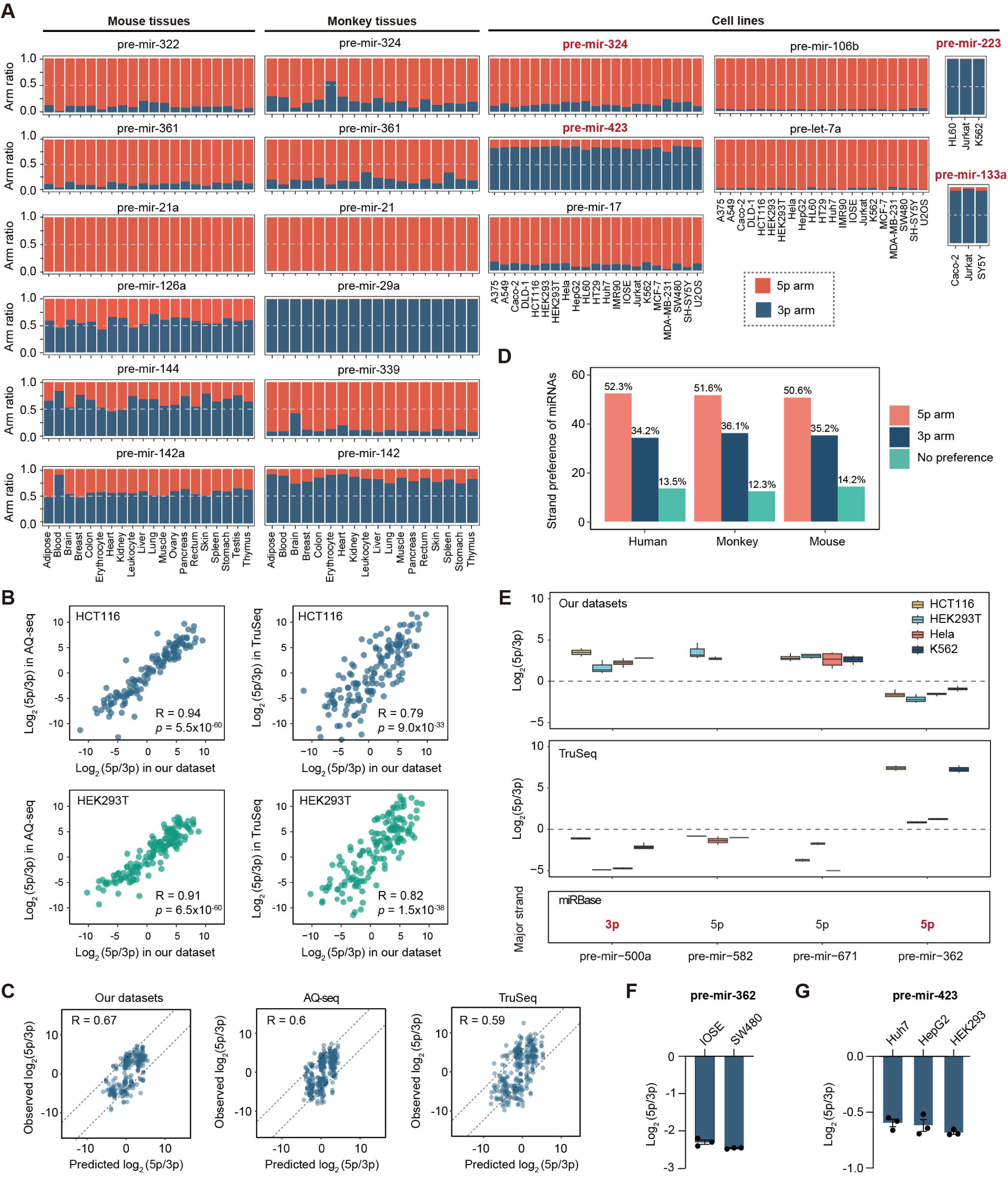
Re-assessment of miRNA strand preference. **(A)** Proportions of the 5’ and 3’ arms of representative miRNAs in various tissues and cell lines, as measured by 4NBoost. miRNAs in red bold indicate that their strand preferences have been previously validated. **(B)** Comparison of 5p/3p ratios for the top 100 miRNAs identified by our datasets, AQ-seq and TruSeq in HCT116 or HEK293T cell lines. Statistical significance was assessed using two-tailed Student’s t-tests. **(C)** Comparison of miRNA strand ratios predicted by the model with those obtained from our datasets (left), AQ-seq (middle), and TruSeq (right) in HEK293T, HCT116, and K562 cells. **(D)** Strand preference of expressed miRNAs across species. miRNAs with over 75% expression from the 5p arm are classified as 5p, those with less than 25% from the 5p arm as 3p, and those in between are categorized as no preference. **(E)** 5p/3p ratios of representative miRNAs across various cell lines measured by our datasets and TruSeq. Annotated strand preferences from miRBase are displayed below, with bold text highlighting annotations that are inconsistent with our datasets. In these plots, the center line represents the median value, the box borders represent the upper and lower quartiles (25th and 75th percentiles, respectively), and the ends of the top and bottom whiskers represent maximum and minimum values, respectively. Biological replicates were as follows: for TruSeq, n = 2 (for HCT116, HEK293T, Hela) and n = 4 (for K562); for 4NBoost, n = 4 (for HCT116, HEK293T, Hela, and K562). **(F-G)** Log_2_-transformed strand ratio of pre-mir-362 (F) and pre-mir-423 (G) obtained from RT-qPCR-based absolute quantification across various cell lines. Each cell line is represented by three biological replicates. Data represent mean ± s.e.m. Source data are provided as a Source Data file.

Examination of strand preference for all detected miRNAs across cell lines and tissues in our 4NBoost data (Supplementary data 4) indicated that >50% of miRNAs preferentially expressed the 5p strand, whereas >30% preferentially expressed the 3p strand, regardless of species (Figure 4D). Additionally, although most miRNAs exhibited consistent strand preferences, a small subset switched the preferred strand depending on the cell type or tissue (Figure 4A). For example, pre-mir-142a produced markedly more 3p than 5p miRNA in mouse blood; but the 5p and 3p strands are present at similar levels in other tissues. Likewise, pre-mir-324 and pre-mir-339 in monkeys, as well as pre-mir-126a and pre-mir-144 in mice, also showed variable strand preferences. Notably, 4NBoost identified several miRNAs with opposite strand selection compared to TruSeq or miRBase annotations, including pre-mir-500a, pre-mir-582, pre-mir-671, and pre-mir-362 (Figure 4E), as well as 15 previously reported miRNAs, such as pre-mir-193a, pre-mir-374a, and pre-mir-454 [29]. Among these, pre-mir-423 and pre-mir-362 were selected, and their strand preference was validated by RT-qPCR. The strand ratios estimated by RT-qPCR were consistent with our dataset but differed from the TruSeq data or miRBase annotations (Figure 4F-G). Taken together, these results indicated that 4NBoost could provide more reliable and accurate miRNA strand preference data than other current methods.

### Re-evaluation of the expression and tissue specificity of miRNA families

miRNA families, which share a common seed sequence and exhibit high sequence similarity, often exert cumulative effects on gene expression [69–73]. However, the extent to which previous biased sequencing approaches affect the accurate measurement of miRNA family expression has not yet been evaluated. We therefore assessed miRNA family-level expression patterns across various cell lines and tissues for comparison with other current miRNA databases.

We observed that miRNA families such as let-7, miR-10, and miR-21 were broadly expressed across human cell lines in 4NBoost data (Figure 5A), which agreed well with previous reports [1, 52, 74, 75]. However, comparison with DIANA-miTED revealed significant discrepancies in certain miRNA family expression patterns compared with 4NBoost data. For example, miR-15, miR-17, miR-19, and miR-29 family expression levels were generally underestimated in DIANA-miTED, while the let-7, miR-10, miR-191, and miR-92 families were overrepresented compared to 4NBoost data (Figure 5B). Further, even more pronounced discrepancies emerged through comparisons of tissue level data between 4NBoost and MirGeneDB (Supplementary Figure 4A-B), which indicated that the miR-126, miR-143, miR-19, miR-23, miR-24, miR-26, and miR-29 families were generally underestimated, whereas let-7, miR-1, miR-103, miR-199, miR-10, miR-181, miR-191 and miR-22 families were overrepresented in MirGeneDB (Supplementary Figure 4C-D). Closer scrutiny of these datasets suggested that these discrepancies likely arose through inaccurate quantification of specific miRNAs, such as miR-19a and miR-19b in the miR-19 family, as well as miR-29a, miR-29b, and miR-29c in the miR-29 family, which led to underestimation of these miRNA families by conventional small RNA sequencing analytical methods (Supplementary Data 3). Consistent with this, previous studies have also shown that miR-29b ranked only 29th in the fixed adapter library derived from DLD-1 cells, whereas it was the most abundant miRNA in the random adapter library [29]. This discrepancy illustrates how miR-29b might be overlooked in colorectal cancer studies. These cumulative results suggested that 4NBoost data could provide a more accurate reference for miRNA family expression, which is essential for elucidating their biological functions and roles in disease processes.

**Figure 5:**
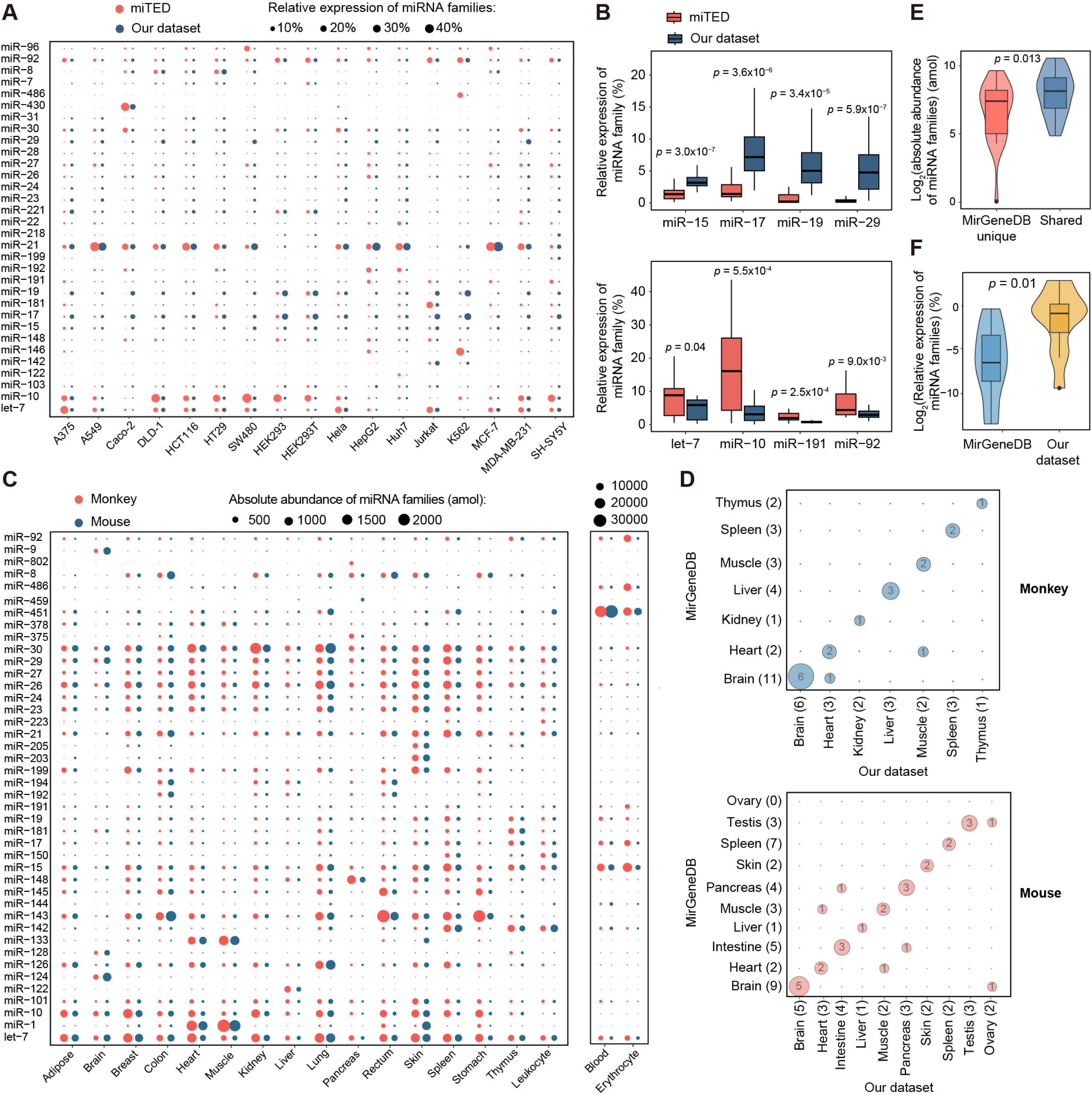
The expression of miRNA families across cell lines and tissues. **(A)** Expression patterns of the top 30 miRNA families in cell lines based on our datasets and the DIANA-miTED database. Circle size represents the proportion of miRNA family expression relative to total miRNA families. **(B)** The representative miRNA families whose expression was underestimated (up) or overrepresented (bottom) in the DIANA-miTED database compared to our datasets. Each box plot summarizes data from the 17 shared cell lines common to both datasets. **(C)** Expression patterns of the top 30 miRNA families in mouse and monkey tissues. Circle size represents the quantification of each miRNA family. Data in (A) and (C) are presented as mean values and their biological replicates: n = 2 (for mouse ovary and testis) or n = 3 (for Caco2, mouse-breast, monkey-rectum, and monkey-stomach) or n = 4 (for the remaining samples). **(D)** Confusion matrix of miRNA families with a tissue-specificity index (TSI) greater than 0.85. Numbers within the bubbles indicate the number of overlapping tissue-specific miRNA families between our datasets and MirGeneDB across various species. Numbers in parentheses on the axes denote the total number of tissue-specific miRNA families identified in each tissue using our datasets (x-axis) and MirGeneDB (y-axis). **(E)** The quantification of tissue-specific miRNA families uniquely identified by MirGeneDB (MirGeneDB unique) or common by MirGeneDB and our datasets (Shared). Each box plot summarizes data from the 10 shared tissues common to both datasets. **(F)** Expression levels of tissue-specific miRNA families uniquely identified by our datasets in the MirGeneDB and our datasets. Each box plot summarizes data from the 10 shared tissues common to both datasets. Statistical significance was assessed using two-tailed Student’s t-tests. Box plots in (B), (E), and (F) depict the median (center line), interquartile range (box), and min-max range (whiskers). Source data are provided as a Source Data file.

We then focused on identifying ubiquitously expressed or tissue-specific miRNA families, which revealed that the let-7, miR-10, miR-23, miR-24, miR-26, miR-27, miR-29, and miR-30 families were highly expressed across all solid tissues, and showed remarkably similar expression levels between mice and monkeys (Figure 5C). These results were in line with the well-documented roles of these miRNA families in various developmental, cellular, and physiological processes essential for most tissues [1]. In contrast with the above broadly expressed miRNAs, we next searched for tissue-specific miRNA families, which are particularly intriguing due to their potential for specialized functions within specific tissues [52, 76]. To assess tissue specificity, we calculated the tissue specificity index (TSI) for miRNA families using a well-established method [77], wherein higher TSI values indicated greater tissue specificity. This analysis uncovered 23 and 30 total tissue-specific miRNA families in 4NBoost data from monkey and mouse tissues, respectively (TSI ≥ 0.85, Table 1). Among these tissue-specific miRNA families, more than 80% were consistent with those identified by MirGeneDB (Figure 5D, Table 2-3), which may be attributable to the high accuracy of relative expression analysis available through all current small RNA sequencing methods [35].

**Table 1.** Tissue-specific miRNA families identified in monkey and mouse tissues using SmRNAQuant datasets.

| Tissues | Monkey | Mouse | Overlap |
| --- | --- | --- | --- |
| Blood | miR-144, miR-451, miR-19, miR-142 | miR-144, miR-451, miR-142, miR-19, miR-15, miR-17 | 4 |
| Brain | miR-124, miR-9, miR-137, miR-132 | miR-124, miR-9, miR-132, miR-137, miR-128, miR-218, miR-154, miR-127 | 4 |
| Erythrocyte | miR-486, miR-92, miR-191, miR-17, miR-15 | miR-486, miR-92 | 2 |
| Heart | miR-499, miR-133, miR-1 | miR-499, miR-133, miR-1 | 3 |
| Leukocyte | miR-150 | miR-150 | 1 |
| Liver | miR-122 | miR-122 | 1 |
| Muscle | miR-133, miR-1 | miR-133, miR-1 | 2 |
| Pancreas | miR-217, miR-216, miR-375 | miR-216, miR-217, miR-459 | 2 |
| Skin | miR-203, miR-205 | miR-203, miR-205 | 2 |
| Colon | - | miR-192, miR-375, miR-194 | 0 |
| Thymus | - | miR-181 | 0 |

**Table 2.** Tissue-specific miRNA families identified in monkey tissues using SmRNAQuant and MirGeneDB.

| Tissues | SmRNAQuant | MirGeneDB | Overlap |
| --- | --- | --- | --- |
| Brain | miR-124, miR-129, miR-132, miR-137, miR-149, miR-9 | miR-124, miR-129, miR-132, miR-137, miR-149, miR-9, miR-127, miR-128, miR-135, miR-138, miR-7, miR-218 | 6 |
| Heart | miR-1, miR-499, miR-135 | miR-1, miR-499 | 2 |
| Kidney | miR-190, miR-204 | miR-190 | 1 |
| Liver | miR-122, miR-192, miR-194 | miR-122, miR-192, miR-194, miR-148 | 3 |
| Muscle | miR-1, miR-133 | miR-1, miR-133, miR-378 | 2 |
| Spleen | miR-150, miR-223, miR-335 | miR-150, miR-223, miR-142 | 2 |
| Thymus | miR-205 | miR-205, miR-155 | 1 |

However, we also found some noteworthy differences with 4NBoost results. First, the tissue-specific miRNA families identified using 4NBoost data were largely subsets of those identified using MirGeneDB (12 out of 17). For example, 6 of 11 brain-specific miRNA families identified by MirGeneDB in monkeys were also detected using 4NBoost. Further analysis revealed that the miRNA families missed by 4NBoost were predominantly expressed at low levels (Figure 5E), and were likely filtered out due to the expression cut-off we applied during the analysis process. Alternatively, 4NBoost analysis identified five tissue-specific miRNA families that were overlooked by MirGeneDB, such as the kidney-specific miR-204 family, heart-specific miR-135 family, and spleen-specific miR-335 family in monkey (Table 2), as well as the ovary-specific miR-135 and miR-202 families in mouse (Table 3). The expression levels of these miRNA families may have been underestimated by conventional sequencing data compared to 4NBoost (Figure 5F), potentially leading to their omission from MirGeneDB. Taken together, the capacity for precise miRNA quantification by 4NBoost facilitates more reliable identification of tissue-specific miRNA families.

**Table 3.**
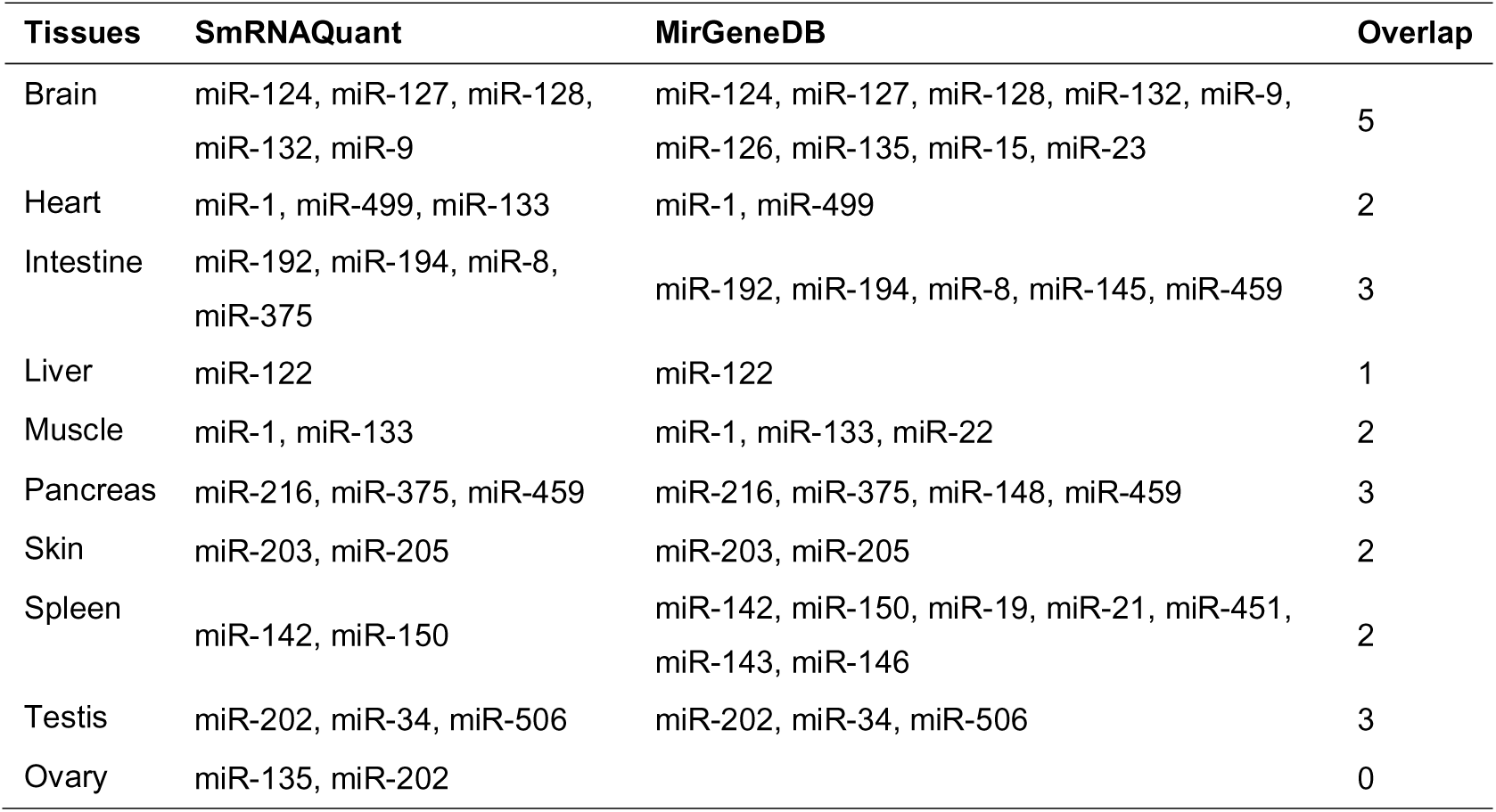
Tissue-specific miRNA families identified in mouse tissues using SmRNAQuant and MirGeneDB.

### Tissue-specific miRNA families are conserved between mice and monkeys

Since tissue-specific miRNAs have been previously shown to exhibit high conservation between mice and humans [77, 78], we next applied 4NBoost to explore the conservation of tissue-specific miRNA families between mice and monkeys. Our results showed that TSI values were quite similar between species for these miRNA families (Pearson’s coefficient = 0.77, Supplementary Figure 5A), which was consistent with their comparable expression profiles. This analysis also showed a high degree of overlap in the tissue-specific expression of miRNA families between the two species (Supplementary Figure 5B, Table 1). For example, we identified several well-documented tissue-specific miRNA families, such as the miR-122 family in liver; miR-9, miR-137, and miR-124 families in brain; miR-133 and miR-1 families in muscle and heart; miR-499 family in heart; miR-144 and miR-451 families in blood; miR-205 family in skin; and miR-216 family in the pancreas [54, 77, 79–82]. Additionally, we identified some previously unreported tissue-specific miRNA families in mice and monkeys, including the miR-203 family in the skin, miR-459 and miR-217 families in the pancreas, as well as miR-19 and miR-142 families in the blood. These findings suggested that tissue-specific miRNA families are conserved across mouse and monkey tissues.

### Re-evaluation of Plant sncRNA abundance

Plant miRNAs and most siRNAs are characterized by 3’-terminal 2’-O-methylation, which substantially impedes adapter ligation and introduces biases in library construction using conventional small RNA sequencing methods. To overcome this limitation, we re-evaluated sncRNA abundance in root, stem, leaf, and flower tissues of the model plant *Arabidopsis thaliana* using 4NBoost. Consistent with the well-established features of plant small RNAs, the 4NBoost profiles were dominated by 21-24 nt sncRNAs, with 21-nt species strongly enriched for 5’ uridine while 24-nt species preferentially carrying adenine (Supplementary Fig. 6A). Compared with the conventional TruSeq method, 4NBoost markedly improved the recovery of 24-nt sncRNAs with a stronger 5’ adenine bias, and significantly enhanced miRNA detection, identifying on average 244 unique miRNAs, nearly 50% more than the 163 miRNAs captured by TruSeq (Supplementary Fig. 6B-D). In addition, TruSeq introduced substantial inaccuracies in miRNA quantification. For instance, miR165a-3p, miR166a-3p, miR168a-3p, miR168a-5p, and miR166e-5p were consistently overestimated, whereas miR161.2, miR172a, miR173-5p, and miR167a-5p were underestimated across all four tissues (Supplementary Fig. 6E). Collectively, these results demonstrate that 4NBoost provides a more accurate and comprehensive representation of plant small RNA abundances, overcoming the limitations of conventional sequencing methods.

### Machine learning-based correction of sequencing bias in sncRNA expression profiles

To enhance the utility of existing sncRNA sequencing datasets affected by library preparation bias, we developed a computational framework that corrects biases in conventional expression profiles, generating quantitatively accurate datasets. We analyzed matched samples processed using two protocols: the widely adopted NEBNext small RNA library preparation and our optimized 4NBoost method. Using these paired datasets, we trained XGBoost regression models to learn and correct library-specific artifacts present in conventional sequencing data (Figure 6A). The correction model exhibited strong performance in reconstructing accurate expression profiles from biased datasets. For NEBNext-prepared libraries, the predicted expression levels showed high concordance with those obtained from 4NBoost, with Pearson correlation coefficients of r = 0.87 in the test set and r = 0.86 in the validation set (Figure 6B). Importantly, the model also improved the ranking accuracy of transcript abundances: the correlation with 4NBoost data increased from r = 0.51 (uncorrected) to r = 0.85 (corrected) in the test set (Figure 6C). Internal validation further confirmed the robustness of the model, yielding similarly high correlations (Pearson’s r = 0.83; Figure 6D). In summary, this regression-based correction framework enables accurate reinterpretation of existing sncRNA-seq datasets by mitigating protocol-induced biases.

**Figure 6:**
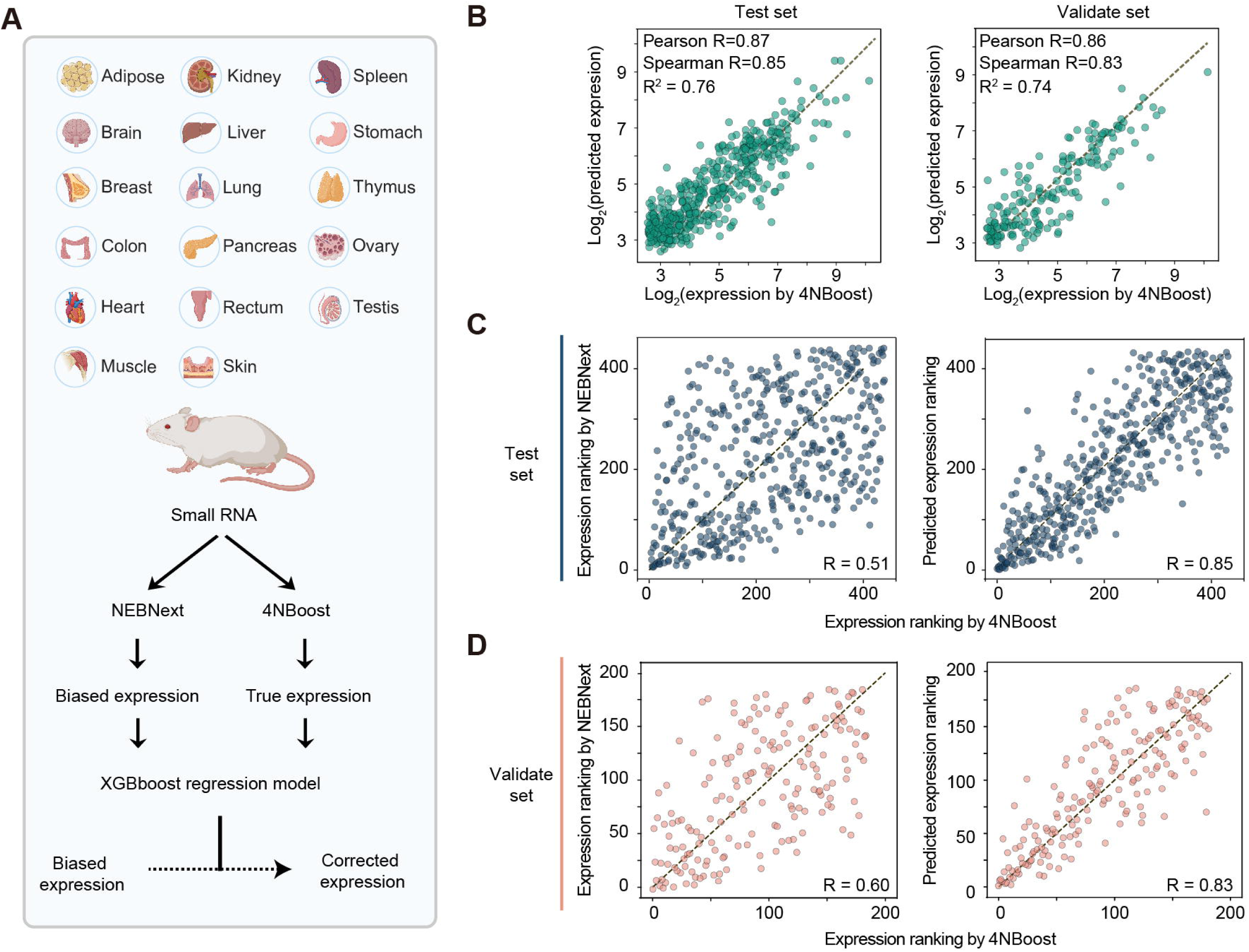
Machine learning-based correction of sequencing bias. **(A)** Schematic workflow of the predictive model, trained on paired datasets generated using NEBNext Small RNA Library Prep (NEBNext) and 4NBoost. This schematic was created with BioRender. **(B)** Concordance between 4NBoost measured and model-predicted miRNA expression values. **(C-D)** Comparison of miRNA expression rankings before (left) and after (right) bias correction, relative to 4NBoost measurements in both test **(C)** and validation sets **(D)**.

### SmRNAQuant: an integrated 4NBoost database

To facilitate access to our 4NBoost data from various cell lines and tissues, we developed SmRNAQuant (Figure 7), an interactive analytical and visualization platform incorporating Django (v4.2.1), Bootstrap (v3.3.7), jQuery (v3.2.1), Python 3.8, and Echarts. This platform enables detailed quantification of small RNAs across different cell lines and tissues, facilitating deeper insights into expression patterns and biological roles. Users can download the complete dataset or generate customized subsets by selecting specific tissues or cell lines of interest. SmRNAQuant also features a “miRNA view” function, which allows users to query individual miRNAs and visualize their expression profiles through bar charts across various tissues and cell types, providing a comprehensive overview of miRNA expression. Additionally, the platform integrates our regression-based correction algorithm, enabling users to calibrate NEBNext-derived datasets and obtain expression values comparable to those generated by 4NBoost. Collectively, SmRNAQuant improves the accessibility and usability of small RNA sequencing data for a broad range of research applications.

**Figure 7:**
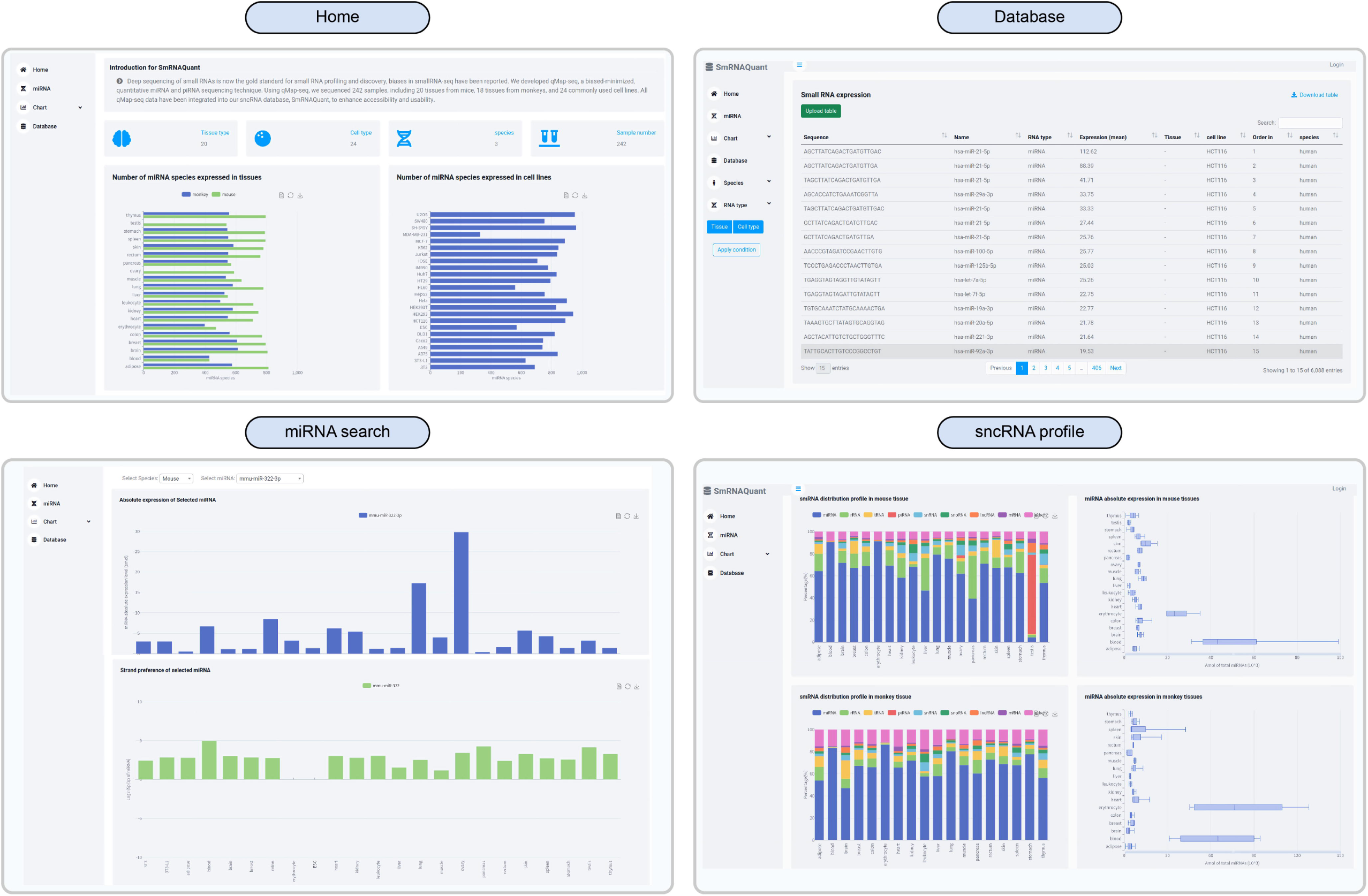
Snapshot depicting the SmRNAQuant interface. Users can view an overview of database statistics from the home interface. The Database interface offers a range of filtering options to refine quantitative information or rankings for sncRNAs of interest, including species, tissue or cell types, and small RNA types. The miRNA search interface provides access to quantitative data and strand preference information for specified miRNAs across various tissues or cell lines. Additionally, the sncRNA profile interface displays the proportions of different sncRNAs expressed in various tissues and cell lines, along with quantitative information for total miRNA.

## DISCUSSION

In this study, we introduce 4NBoost, a bias-minimized and quantitatively calibrated method for estimating absolute sncRNA profiling. We also introduce SmRNAQuant, a web-based database designed to host quantification datasets of sncRNAs. To further enhance the usability of 4NBoost and SmRNAQuant, we developed a machine learning-based correction model to adjust for biases inherent in conventional sncRNA sequencing data. Together, our work provides a large-scale and systematic atlas of quantification of sncRNAs across a diverse range of mammalian tissues and widely used cell lines.

Compared with external databases, our datasets provide more accurate small RNA information. First, it offers a more precise ranking of miRNAs in tissues and cell lines, which is crucial for researchers to select research targets. For example, our results indicate that, compared to existing databases, the expression levels of the miR-29 and miR-19 families are significantly higher, whereas those of the miR-10 and let-7 families are significantly lower. These differences are mainly attributed to the structural preferences of the T4 ligation enzyme. Second, our datasets provide more reliable information on miRNA strand preference, which is consistent with both experimental and model-based findings reported in the literature. Moreover, in comparison to TruSeq sequencing data and miRBase, we identified strand annotation deviations in certain miRNAs, such as pre-mir-500a and pre-mir-362, which could greatly affect the prediction of their target genes. Third, our datasets provide a higher signal-to-noise ratio for tissue-specific miRNAs, making it a more reliable dataset for studying miRNA conservation across species.

Previous studies demonstrated that miRNA activity depends on reaching a threshold level of expression [44], with only the most abundant miRNAs in a cell capable of mediating effective target suppression [43]. A recent study further showed that mRNA stability begins to decrease when miRNA abundance reaches approximately 1,000 miRNA transcripts per million (TPM) [40]. In mouse oocytes, rapid growth and dilution of miRNAs during development were found to lower their effective concentrations, producing a knockdown-like effect insufficient for robust repression [42]. Consistently, silencing activity of miRNAs was maintained at 1.5 nM but lost at 0.3 nM. Collectively, these findings highlight the tight coupling between intracellular miRNA concentration and functional activity. However, absolute expression levels have so far been determined for only a subset of miRNAs in limited cell and tissue types [43, 45–47]. In this context, our datasets provide a more comprehensive and systematic reference, enabling the prediction of which miRNAs are likely to be functionally active and guiding the selection of the most relevant candidates for downstream studies.

In addition, we developed an XGBoost-based correction framework that converts conventional small RNA-seq data into quantitatively accurate expression profiles. By eliminating the need for re-sequencing, it substantially expands the value of existing data and facilitates more reliable downstream analyses. Leveraging the unbiased quantification provided by 4NBoost, SmRNAQuant integrates this correction model to serve as a more reliable and precise reference resource for small RNA research.

Despite its strengths, this study has several limitations. First, the effectiveness of 4NBoost for constructing sncRNA libraries depends on a minimum input of 1 ng of total RNA. Below this threshold, data quality declines markedly, primarily due to incomplete removal of excess 3’ adapters, which leads to the accumulation of adapter–adapter PCR artifacts [56]. Future optimization efforts may focus on enhancing 3’ adapter removal efficiency or exploring alternative strategies, such as using a cocktail of RNA ligases to reduce ligation bias and potentially eliminate the need for terminal degenerate sequences. It should also be noted that although 4NBoost reduces ligation bias and improves quantification accuracy, the method still exhibits some bias and therefore cannot guarantee fully precise absolute quantification. Second, 4NBoost is specifically designed for quantifying sncRNAs that share compatible chemical termini (5’ monophosphate and 3’-OH or 3’-O-methyl ends), such as miRNAs, siRNAs, and piRNAs. However, other classes of sncRNAs, including small nucleolar RNAs (snoRNAs), small nuclear RNAs (snRNAs), tRNA-derived small RNAs (tsRNAs), and rRNA-derived small RNAs (rsRNAs), often contain noncanonical 5’ or 3’ ends or internal modifications that interfere with ligation and reverse transcription in the current 4NBoost protocol. Several specialized methods have been developed to address these challenges [83–86], and integrating such techniques with 4NBoost may broaden its scope. Finally, the limited diversity of samples included in our current database may restrict the generalizability of our findings and reduce the applicability of the data across broader biological contexts. To overcome this limitation, future studies should apply 4NBoost to a wider range of tissues, developmental stages, and disease states. In parallel, our machine learning–based correction framework offers a complementary solution by enabling integration and bias correction of existing sncRNA-seq datasets. Together, these efforts will expand sample representation and enhance the robustness and translational potential of the resulting small RNA expression database.

## METHODS

### Cell culture and RNA extraction

All cell lines used in this study were purchased from the American Type Culture Collection (ATCC), National Collection of Authenticated Cell Cultures (NCACC), or Shanghai Chuanqiu Biological Technology Co., Ltd (SCBT) (Supplementary Data 1) and were authenticated by the respective providers. Cell lines were revived and cultured in a 6 cm dish at 37°C with 5% CO_2_. Once the cells reached 90% confluence, they were collected with 1 mL of TRIzol reagent (Invitrogen). Next, 200 µL of trichloromethane was added and mixed thoroughly. The solution was then centrifuged at 12,000 g for 15 minutes at 4°C. The upper aqueous phase (∼450 µL) was transferred into a new 1.5 mL EP tube, followed by the addition of 900 µL of ethanol. The solution was mixed by inversion, left at -30°C for 30 minutes, and then centrifuged again at 12,000 g for 15 minutes at 4°C. After removing the supernatant, the pellet was washed twice with 80% ethanol, air-dried for 2-3 minutes at room temperature, and dissolved in DEPC-treated water. RNA concentration was measured using a Nanodrop spectrophotometer, and RNA quality was assessed using the Agilent Bioanalyzer 2200. Each RNA sample was diluted to a final concentration of 100 ng/µL and stored at -80°C until use.

### Tissue collection and RNA extraction

This study was conducted using both male and female mice and Crab-eating monkeys. 4 weeks old male and female BALB/c mice were purchased from Shanghai Lingchang Biotechnology Co., Ltd, and animal protocol was approved by the Center for Excellence in Molecular Cell Science, Chinese Academy of Sciences (SIBCBS2182005007). 4 years old male and female Crab-eating monkey tissues were generously provided by the Jinyi Chen’ lab at Shanghai Institute of Materia Medica, Chinese Academy of Sciences.

Fresh blood samples were collected into EDTA anticoagulant tubes and diluted with PBS at a 1:1 ratio. 3mL of separation solution (Solarbio) and 3 mL of the diluted blood were added to a 15 mL centrifuge tube and centrifuged at 800 g for 30 minutes at room temperature, resulting in distinct layers containing plasma, white blood cells, separation solution, and red blood cells. The red blood cell layer was carefully transferred into a new 1.5 mL EP tube, mixed with 1 mL of Trizol, and stored at -20°C until RNA extraction. The white blood cell layer was transferred into a new 15 mL centrifuge tube, washed with 10 mL of PBS, and centrifuged at 250 g for 10 minutes at room temperature. After centrifugation, the supernatant was removed, and the precipitate was resuspended with red blood cell lysis buffer, then centrifuged at 250 g for another 10 minutes. The supernatant was discarded, and the precipitate was resuspended with 1 mL Trizol. After thorough mixing, the sample was stored at -20°C until RNA extraction.

Mouse and monkey lung, breast, stomach, colon, rectum, kidney, brain, skin, liver, ovary, testis, fat, pancreas, heart, thymus, muscle, and spleen tissues were collected, cut into small pieces, and submerged in an appropriate amount of RNAlater (Thermo). After being stored overnight at 4°C, the samples were transferred to -20°C. For RNA extraction, 50-100 mg of tissue was ground using a cryogenic grinder or a mortar (see Supplementary Table 2 for the specific grinding method for each tissue), and RNA was purified as described above.

### Small RNA spike-in design

We randomly generated 1,000 sequences with lengths ranging from 20 to 30 nt and predicted their secondary structures using RNAfold. Sequences were selected based on the following criteria: GC content between 40% and 60%, a predicted secondary structure with a ΔG greater than -3, and no alignment with common model organism genomes. Finally, 23 sequences were chosen as spike-in candidates. To evaluate the ligation efficiency of RNAs with 2’-Ome at the 3’ ends, seven of these candidates were randomly selected. The 2’-OH at their 3’ end was replaced by 2’-Ome, and one of their bases was altered to distinguish them from their 2’-OH counterparts. All spike-ins were synthesized by Suzhou Olipharma Co., Ltd.

### Small RNA library preparation

One hundred nanograms of total RNA with 0.05 µL of 20 nM EM or RM spike-in control oligos was ligated to a 0.55 µM 3’ randomized adapter using 25 U/µL T4 RNA ligase 2 truncated KQ (NEB) in 0.83X T4 RNA ligase reaction buffer (NEB), supplemented with 20% PEG 8000 (NEB) at 25°C overnight. The ligated RNA was then annealed with 1.28 µM RTP. To remove excess 3’ adapters, 3.33 U/µL 5’ deadenylase (NEB) and 0.2 U/µL lambda exonuclease (NEB) were added, followed by incubation at 30°C for 1 hour and 37°C for 2 hours, respectively. Subsequently, the product was ligated to a 0.55 µM 5’ randomized adapter using 0.4 U/µL of T4 RNA ligase 1 (NEB) in 0.74X T4 RNA ligase reaction buffer, supplemented with 0.7 mM ATP and 17.3% PEG 8000 (NEB) at 25°C for 1 hour. Reverse transcription was carried out using 5 U/µL of ProtoScriptII reverse transcriptase (NEB) in 1X ProtoScriptII RT reaction buffer (NEB), containing 0.5 mM dNTPs (Promega) and 10 mM DTT (NEB), at 50°C for 1 hour. The resulting cDNA was then amplified using 0.02 U/µL of KOD-Plus-Neo (TOYOBO) in 1X PCR Buffer, along with 0.5 µM of both forward and reverse primers, 1.5 mM MgSO_4_ (TOYOBO), and 0.2 mM dNTPs (TOYOBO). After PCR amplification, the cDNA was gel-purified on a 6% polyacrylamide gel to remove adapter dimers and sequenced using the Illumina NovaSeq6000 sequencing system. The sequences of adapters, along with the sequences and specific mixing ratios of EM or RM spike-ins, are provided in Supplementary Table 3. Mouse tissue miRNA libraries (excluding blood, erythrocytes, and leukocytes) were also prepared using the NEBNext^®^ Multiplex Small RNA Library Prep Set for Illumina^®^, with 100 ng of total RNA as input.

### Quantitative real-time PCR

We selected miR-221-3p (whose abundance spanned the entire linear range of our spike-in standard curve) and miR-21-5p (the most abundant in nearly all cell lines) for evaluation of quantification accuracy at different concentrations. The reverse transcriptase reactions consisted of 100 ng of purified total RNA, 50 nM stem-loop RT primer, 1X reverse transcriptase M-MLV buffer (TAKARA), 0.25 mM of each dNTP (Promega), 10 U/µL of reverse transcriptase M-MLV RNase H- (TAKARA), and 1.07 U/µL of RNase inhibitor (Thermo). The total volume of the reactions was 7.5 µl, and they were incubated in a Bio-Rad T100 Thermal Cycler under the following conditions: 30 minutes at 25°C, 1 hour at 42°C, and 15 minutes at 75°C, before being held at 4°C. RT products were diluted five times with deionized water. Each real-time PCR assay for the microRNA (10 µL volume) contains 2 µL of diluted RT product, 5 µL of 2X Taq Pro Universal SYBR qPCR Master Mix (Vazyme), and 1.5 µM of both forward and reverse primers. The reactions were carried out in an Applied Biosystems® QuantStudio™ 6 Flex Real-Time PCR System in 384-well plates, with the following cycling conditions: an initial denaturation at 95°C for 30 seconds, followed by 40 cycles of denaturation at 95°C for 10 seconds and annealing/extension at 60°C for 30 seconds. The sequences of RT and PCR primers are listed in Supplementary Table 4.

To validate the ratio of miRNA-5p and -3p strands, hsa-mir-326 and hsa-mir-423 were selected. Reverse transcription (RT) and dilution steps were performed as described previously. Each real-time PCR reaction was carried out in a 10 µL volume containing 2 µL of diluted RT product, 5 µL of 2× Taq Pro HS U+ Probe Master Mix (Vazyme), 1× ROX Reference Dye2 (Vazyme), 0.1 µM TaqMan probe, and 0.2 µM each of the forward and reverse primers. Amplifications were performed on an Applied Biosystems® QuantStudio™ 6 Flex Real-Time PCR System using 384-well plates under the following cycling conditions: 37 °C for 2 minutes and 95 °C for 10 seconds, followed by 45 cycles of 95 °C for 10 seconds and 60 °C for 30 seconds. The sequences of the RT and PCR primers are provided in Supplementary Table 4.

### Analysis of small RNA sequencing

Raw data (read1) were pre-processed using the FASTX Toolkit (v0.0.14). The first 70 bp of each read were retained, and low-quality sequences were filtered out (-q 20, -p 90). After quality filtering, adapter sequences were trimmed, requiring a minimum 10-nucleotide match at the 3’ end. Reads without a detected adapter or shorter than 32 bp were discarded. Only reads with “CGA” at positions 4-6 were retained, corresponding to the 5’ random adapter. UMI extraction was performed with UMI_tools (v0.5.1) using the following parameters: extract --extract-method regex --bc-pattern=”^(?P<UMI_1>.[3])(?P<CELL_1>.[3])(?P<UMI_2>.[4])([A,T,C,G][5,100])(?P <UMI_3>.[4])$”. Processed reads were aligned to the target genome using Bowtie (v1.2.1.1) with parameters --best -v 0. Duplicate reads were removed with UMI_tools using the following parameters: dedup --edit-distance-threshold 1 --soft-clip-threshold 0 --method adjacency. Unaligned reads were mapped to the spike-in reference sequences using Bowtie with parameters --best -v 1 --norc and de-duplicated in the same manner. The remaining unmapped reads were aligned to the isomiR reference library with the same parameters and de-duplicated. Molecules aligned to the genome were merged with isomiR molecules for downstream analysis. Molecules were annotated based on the following priority: mature miRNA > isomiR > tRNA > rRNA > snoRNA > snRNA > scRNA > scaRNA > YRNA > lncRNA > mRNA > piRNA > novel miRNA. Mature miRNA, isomiR, piRNA, and novel miRNA were annotated based on exact sequence and length matches, while other small RNA categories were aligned to known sequences using Bowtie with parameters -k=100 -v=0 -norc. Reads that aligned to the genome but lacked known annotations were classified as “others”. The expression of molecules in each library was normalized using the standard curve generated from the external spike-ins.

### Prediction of novel miRNAs

To identify potential novel miRNAs in our library, we applied miRDeep2 (v0.1.2) to reads that did not align to known annotations. Novel miRNAs predicted by miRDeep2 were further filtered based on the following criteria: (1) no match to any known sequences in the Rfam database; (2) a total of at least 10 reads supporting the miRNA in the library; and (3) the mature miRNA sequence must have a corresponding star sequence. Only miRNAs that satisfied all three conditions were deemed reliable, and their precursor miRNA, mature miRNA, and star sequences were extracted. Additionally, if miRge3.0 predicted pre-miRNA sequences with expression from both strands, and the 3p or 5p strand had more than 10 reads and was not identified by miRDeep2, those sequences were incorporated into the miRDeep2 prediction results. Together, these sequences constituted the reference set for novel miRNAs.

### Expression of miRNA families

To assess the expression patterns of miRNA families, we extracted all miRNA family annotations from the latest version of MirGeneDB (v2.1). For each miRNA precursor, all mature forms have been considered as family members. To reduce potential bias from multiple precursors, duplicated mature miRNAs (e.g., those derived from different precursors within the same family) were counted only once.

### Prediction of miRNA arm ratio

We analyzed the miRNA arm ratio as previously described [68] with the following model:

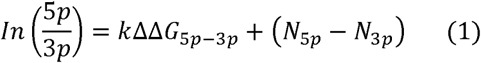

Where *k* and *N*_5p(3p)_ represent the constant for the relative thermodynamic stability and the constant corresponding to the 5′ end identity, respectively.

### Tissue specificity index

To compute the tissue specificity index, we used the formula described previously [77]. The same formula was applied to cells to provide a cell specificity index. Specifically, the TSI for a miRNA j is calculated as

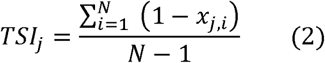

Where *N* is the total number of tissues measured and *x_j,i_* is the expression score of tissue *i* normalized by the maximal expression of any tissue for miRNA *j*.

### Model development and validation

To enhance model compatibility, we integrated both in-house data generated using NEB kits and an external raw dataset prepared with the same kits. After preprocessing, the datasets were merged. To minimize background noise, miRNAs with expression levels below 5 amol in the 4NBoost data were excluded. The filtered dataset was split into three subsets: 72% for training, 20% for testing, and 8% as an independent validation set not involved in model training. In addition to NEB-biased expression levels as core input features, we extracted several parameters as auxiliary features, including (i) Sequence features: RNA length, nucleotide composition (A/T/C/G ratio), and GC content; (ii) Thermodynamic parameters: minimum free energy, free energy difference between 5’ and 3’ termini, and total duplex free energy; (iii) Terminal motifs: the first 1, 3, 4, and 6 nucleotides at both 5’ and 3’ ends; (iv) miRNA structural features: paired ratio, bulge density, internal loop density, hairpin density, and pairing continuity. Feature correlation analysis showed that NEB-biased expression contributed most significantly to the prediction performance (R = 0.614). Terminal sequence features and some thermodynamic parameters (delta_G_sum and MFE) showed moderate correlation (R > 0.14), whereas structure-related features contributed minimally (R < 0.05). Therefore, the five miRNA-structure-related features were excluded from the final feature set.

Using default hyperparameters, we benchmarked four tree-based algorithms: Random Forest (RF), Gradient Boosting Decision Tree (GBDT), LightGBM, and XGBoost. Comparative performance analysis identified XGBoost as the best-performing framework for our dataset, a robust machine learning method known for its robustness and high predictive accuracy. Model training was conducted on the designated training dataset, with hyperparameter optimization performed using the Hyperopt package in Python.

The final model architecture incorporated the following optimized parameters: a learning rate (eta) of 0.4, num_boost_round of 10, max_depth of 8, min_child_weight of 14, subsample of 0.75, colsample_bynode of 0.5, colsample_bytree of 1.0, and reg_lambda of 4.0. Model performance was evaluated on both a test set and an independent external validation dataset. Two primary evaluation metrics were used: the coefficient of determination (R²) to quantify the variance explained by the model in predicting absolute expression values, and Spearman’s rank correlation coefficient to measure the concordance between predicted and actual expression rankings.

### Download of the genome sequence and small RNA annotation

The reference genomes for mouse (mm10), human (hg38), and crab-eating monkey (macFas5) were downloaded from the UCSC Genome Browser (https://hgdownload.soe.ucsc.edu/). Genome annotations for crab-eating monkeys were obtained from UCSC, while annotations for the mouse and human genomes were downloaded from Gencode (https://www.gencodegenes.org/). Small RNA annotations for humans and mice were sourced as follows: miRNA annotations were downloaded from miRBase (release 22, https://mirbase.org/download/) and retained only unique sequences. tRNA annotations were obtained from Ensembl (release 110, https://www.ensembl.org/biomart/martview/) and GtRNAdb (release 21, http://gtrnadb.ucsc.edu/index.html), with duplicate entries removed. rRNA annotations were retrieved from Ensembl and NCBI (https://www.ncbi.nlm.nih.gov/nucleotide/), and duplicates were similarly removed. piRNA annotations were sourced from the piRBase gold standard set (release v3.0, http://bigdata.ibp.ac.cn/piRBase/). Annotations for snRNA, scRNA, snoRNA, scaRNA, YRNA, mRNA, and lincRNA were downloaded from Ensembl. Small RNA annotations for crab-eating monkeys were sourced as follows: miRNA annotations from miRBase for rhesus macaques, with unique sequences retained. Annotations for rRNA, tRNA, snRNA, and scRNA from UCSC (https://hgdownload.soe.ucsc.edu/goldenPath/macFas5/database/rmsk.txt). Additionally, to comprehensively classify small RNAs derived from functional RNAs, we combined the annotated functional RNA sequences from the rhesus macaque genome with corresponding annotations from the crab-eating monkey genome. Annotations for rhesus macaques were retrieved from NCBI (https://www.ncbi.nlm.nih.gov/).

### Web server and database implementation

To create an efficient and engaging web interface for the SmRNAQuant database, we employed the following tools: Django (v4.2.1) for web development, Bootstrap (v3.3.7) for responsive design, jQuery (v3.2.1) for dynamic interactions, Python 3.8 for backend processing, and ECharts for interactive data visualization. Data management and retrieval were conducted using MySQL (v8.0.35).

## DATA AVAILABILITY

The deep-sequencing data have been deposited at the National Center for Biotechnology Information NCBI Gene Expression Omnibus (http://www.ncbi.nlm.nih.gov/geo/) database under accession number GSE279145. The following publicly available datasets were used: K562 AQ-seq data from GSE158025; K562 Truseq data from GSE102497; HCT116 AQ-seq data from GSE230544; HCT116 Truseq data from GSE180613 and GSE189908; HEK293T and Hela AQ-seq data from GSE123627; HEK293T and Hela Truseq data from GSE57295; HEK293T IsoSeek data from PRJNA867189; mouse liver and brain Truseq data from GSE227578; Mouse kidney NEBNext data from PRJNA759746; Mouse liver NEBNet data from GSE123346. Source Data are provided with this paper.

## CODE AVAILABILITY

There was no new code generated in this study. All code or software used is publicly available and has been specified in the Method section.

## ACKNOWLEDGEMENTS

We are grateful to Jin Ren and Xinming Qi for providing Crab-eating monkey tissues and Jun Zhu for providing *Arabidopsis thaliana* tissues. We thank all members of L. Wu’s laboratory for their discussion and comments on this project; Y. Xu for his assistance with high-performance computing; and the staff at the HPC storage and network service platform of SIBCB for supplying the computing resources. Schematics in Figs. 2a and 6a were created using BioRender.com and are included under a paid publication license obtained from BioRender. We acknowledge BioRender for providing the platform and tools for scientific illustration. This work was supported by the National Key R&D Program of China (2022YFA1303301), the Strategic Priority Research Program of the Chinese Academy of Sciences (XDB0570000), and the National Key R&D Program of China (2021YFA1100201) to Ligang Wu; the National Natural Science Foundation of China (32200696) and the Youth Innovation Promotion Association CAS to H.Z.

## AUTHOR CONTRIBUTIONS STATEMENT

L.W. and H.Z. conceived and designed the study. Y.Z. and H.Z. conceived and developed the methodology of 4NBoost. B.X., R.Z., W.X., and Y.Z. collected the cells and tissues. B.X., R.Z., and Y.Z. isolated all the RNAs. Y.Z. generated all sncRNA-seq libraries. W.X. designed spike-ins and performed computational analyses. W.X., H.Z., and Y.Z. interpreted the data. W.X., H.Z., L.W., and Y.Z. wrote the manuscript.

## COMPETING INTERRESTS STATEMENT

The authors declare no competing interests.

---

**Supplementary Figure 1:**
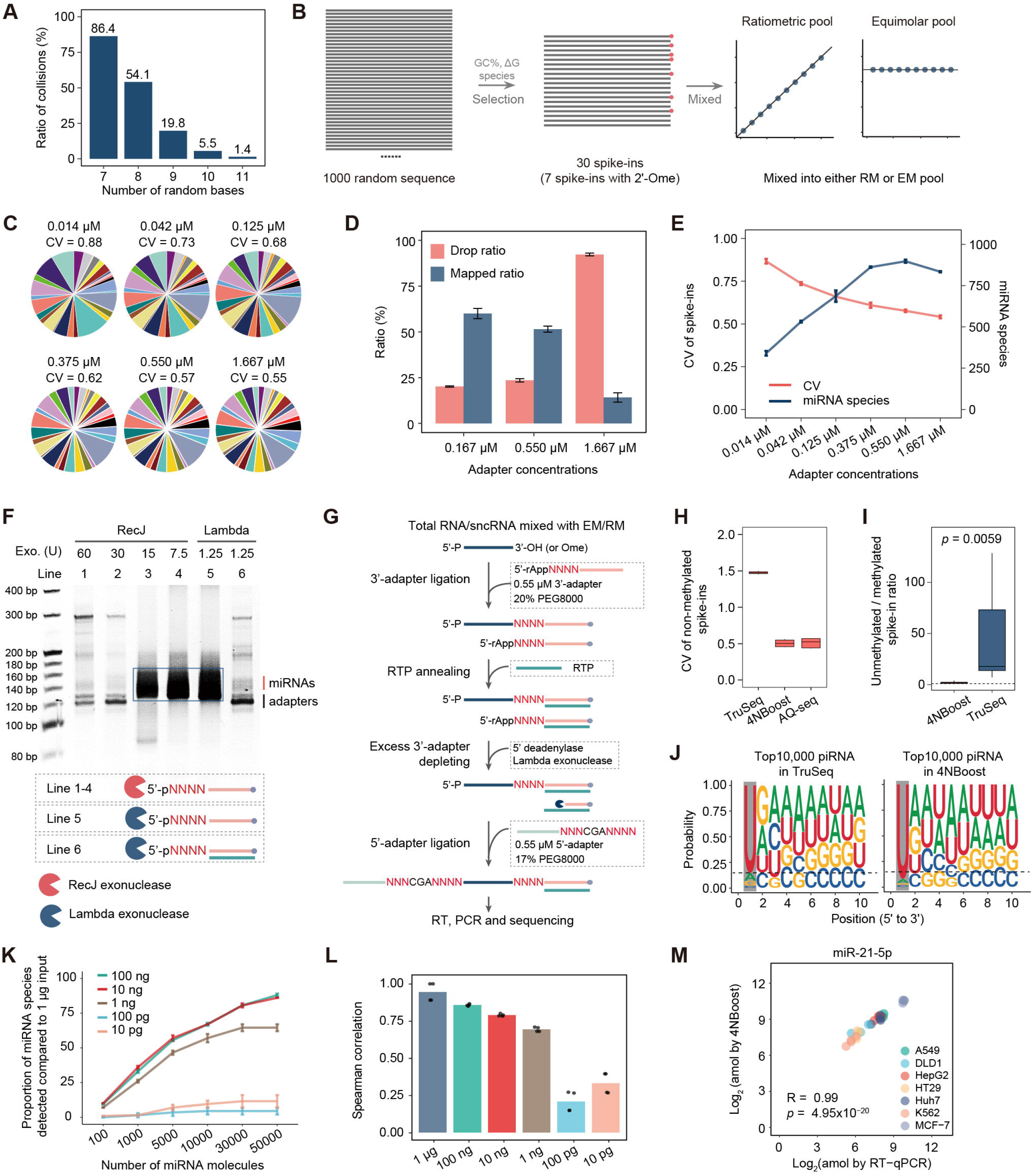
Establishment of the 4NBoost method. **(A)** Histogram illustrates the collision ratios predicted by the AmpUMI model for UMIs with varying random base lengths. The x-axis indicates the UMI random base lengths, while the y-axis displays the predicted collision ratios. **(B)** Schematic illustrating the design, selection, and pooling process for equimolar and ratiometric spike-ins. **(C)** Pie charts showing the proportion of 30 equimolar spike-ins detected under different concentrations of 3’ and 5’ adapters. The coefficient of variation (CV) is shown as the mean value. **(D)** Histogram showing the dropping and mapping ratios of libraries at varying concentrations of adapters. **(E)** Dual line graph illustrating the CVs of equimolar spike-ins and the number of detected miRNA species across different adapter concentrations. The x-axis represents adapter concentrations, while the left y-axis displays the CVs of equimolar spike-ins, and the right y-axis presents the number of detected miRNA species. **(F)** Comparison of the efficiency of Lambda exonuclease and RecJ exonuclease in depleting excess 3’ adapters. In a typical reaction, more than 30 U RecJ or 1.25 U Lambda exonuclease can efficiently remove excess 3’ adapter, resulting in the successful miRNA library construction (lines 1, 2, and 6). However, if the concentration of RecJ exonuclease is below 30 U, excess 3’ adapters can ligate to 5’ adapters, resulting in a large amount of byproducts (lines 3 and 4, highlighted by the blue frame). Additionally, the high efficiency of Lambda exonuclease in depleting 3’ adapters requires the annealing of 3’ adapters and RTP (lines 5 vs. line 6). **(G)** Schematic representation of the 4NBoost library preparation workflow. **(H)** Box plots of CVs for equimolar spike-ins distribution using total cellular RNA as background. Center line represents the median; boxes indicate 25th–75th percentiles; whiskers show minimum and maximum values. The biological replicates for these experiments were n = 2 for TruSeq, n = 4 for 4NBoost, and n = 10 for AQ-seq. **(I)** Relative abundance of non-methylated spike-ins compared to their terminal methylated counterparts. Seven spike-in pairs from equimolar spike-in pools were spiked into 100 ng of HEK293 total RNA. Each pair consisted of one non-methylated and one terminally methylated RNA molecule with nearly identical sequences, differing by only two nucleotides. Higher relative abundance indicates more efficient ligation of the non-methylated spike-in compared to its methylated counterpart. The biological replicates for these experiments were n = 2 for TruSeq and 4NBoost. Box plots depict the median (center line), interquartile range (box), and min-max range (whiskers). The dashed line is the y-axis at 1. **(J)** Relative nucleotide composition of the top 10,000 piRNAs in TruSeq and 4NBoost libraries from mouse testis. **(K)** Proportion of miRNA species detected from libraries at different dilutions, compared to the miRNA species detected from 1 µg of input RNA. **(L)** Spearman correlation of miRNA expression profiles from libraries at different dilutions, compared to the profile from 1 µg of input RNA. **(M)** Scatter plot illustrating the correlation between the predicted miR-21-5p abundances measured by 4NBoost and RT-qPCR across multiple cell lines. The number of biological replicates per cell line was three for DLD1 and HT29 and four for all other cell lines. All p-values were calculated using two-tailed Student’s t-tests. Data in (C), (D), (E), (K), and (L) represent two biological replicates. Data in (D), (E), (K), and (L) represent mean ± s.e.m. Source data are provided as a Source Data file.

**Supplementary Figure 2:**
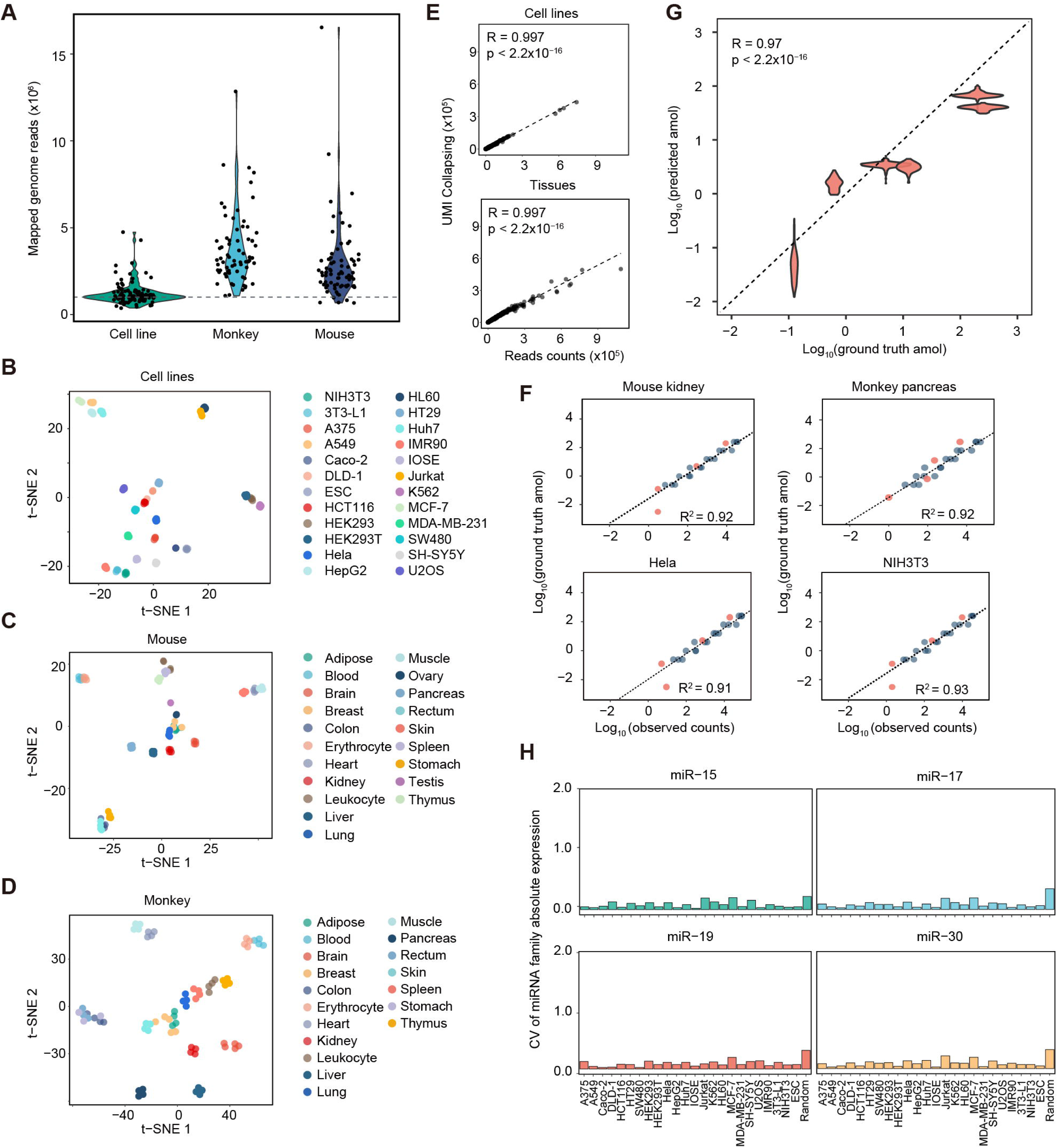
The data quality of 4NBoost libraries from cell lines and tissues. **(A)** Violin plot showing the distribution of mapped reads across all small RNA libraries. **(B-D)** t-SNE projection based on miRNA expression in cell lines**(B)**, mouse (**C**), and monkey (**D**) tissues, with each cell line or tissue represented by three to four biological replicates. **(E)** Scatter plot comparing total read counts and UMI-deduplicated unique molecule counts for the top 100 most abundant miRNAs of each sample. Each point represents one single miRNA within an individual sample of a cell line or tissue. **(F)** Represented examples demonstrating the quantification precision achieved by 4NBoost. The standard curve for each library was established using ratiometric spike-ins with known absolute molecule numbers. Four additional spike-ins, each with varying but known absolute molecule counts, were plotted to evaluate the precision of these standard curves. The x-axis represents the sequencing molecule numbers of each spike-in, and the y-axis shows their corresponding absolute molecule numbers. The R^2^ values indicate the fit of these standard curves. Blue points represent the 30 ratiometric spike-ins, and red points represent the four additional spike-ins. **(G)** Assessment of 4NBoost’s quantification accuracy. Predicted molecule counts of four spike-ins in each sample were calculated from standard curves and plotted against their actual molecule counts. The strong correlation between predicted and ground truth values demonstrates that 4NBoost can accurately quantify the molecule counts of sncRNAs. **(H)** Assessment of miRNA family expression stability within each cell line. Bars indicate the mean CV of representative miRNA family expression levels across biological replicates. Biological replicates: n = 3 (Caco2) or n = 4 (remaining samples). All p-values were calculated using two-tailed Student’s t-tests.

**Supplementary Figure 3:**
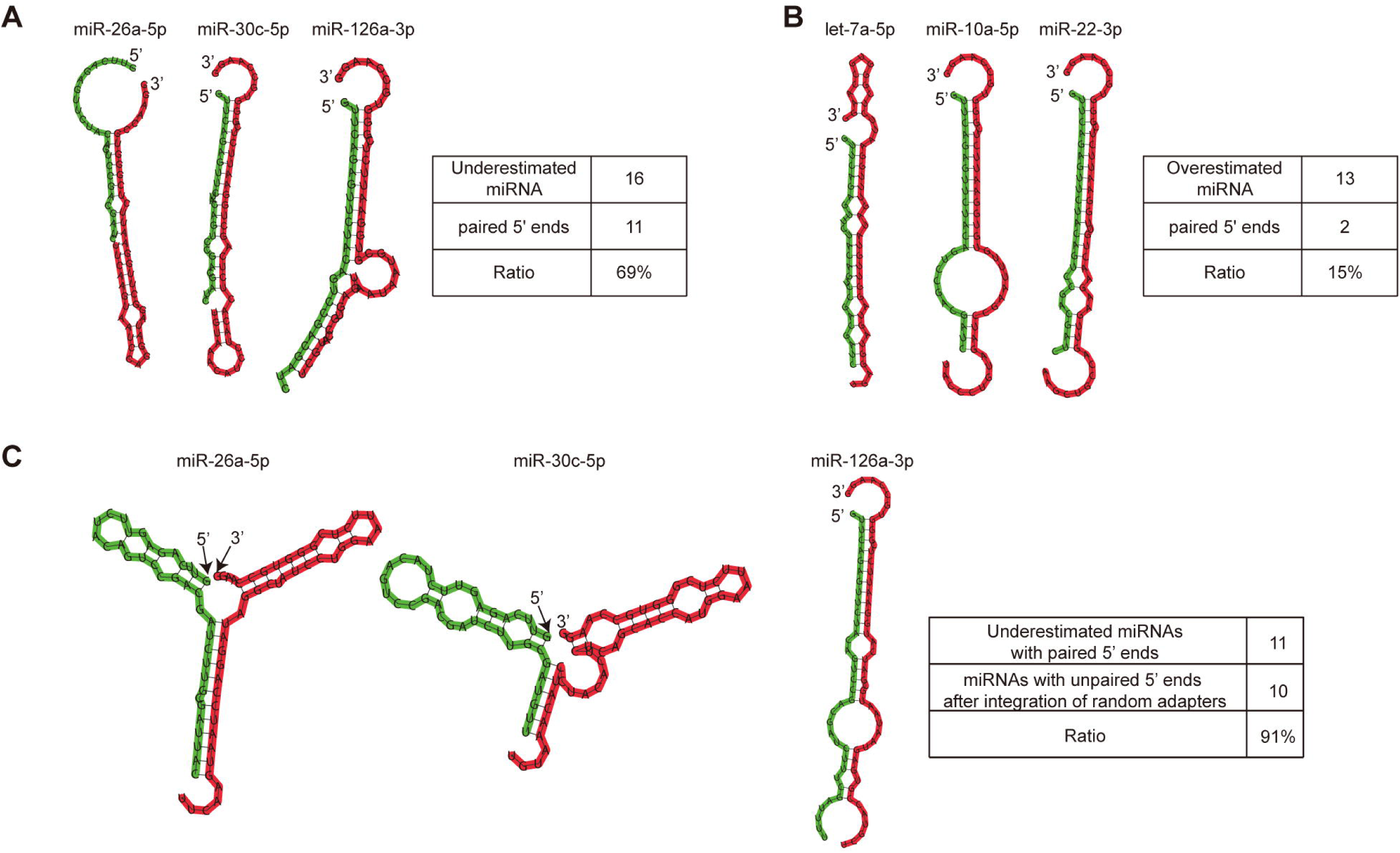
Predicted heterodimer structures formed between miRNAs and fixed or random 5’ adapters (A-B) Predicted heterodimer structures formed between underestimated **(A)** or overestimated miRNAs **(B)** and fixed 5’ RNA adapters used in the TruSeq protocol. **(C)** Predicted heterodimer structures formed between these underestimated miRNAs, as shown in (A), and random 5’ adapters used in the 4NBoost protocol. Three representative structures are shown on the left. Sequences with a red background indicate the miRNAs, while sequences with a green background represent the 5’ RNA adapters. Statistical results for all miRNAs are summarized in the table on the right.

**Supplementary Figure 4:**
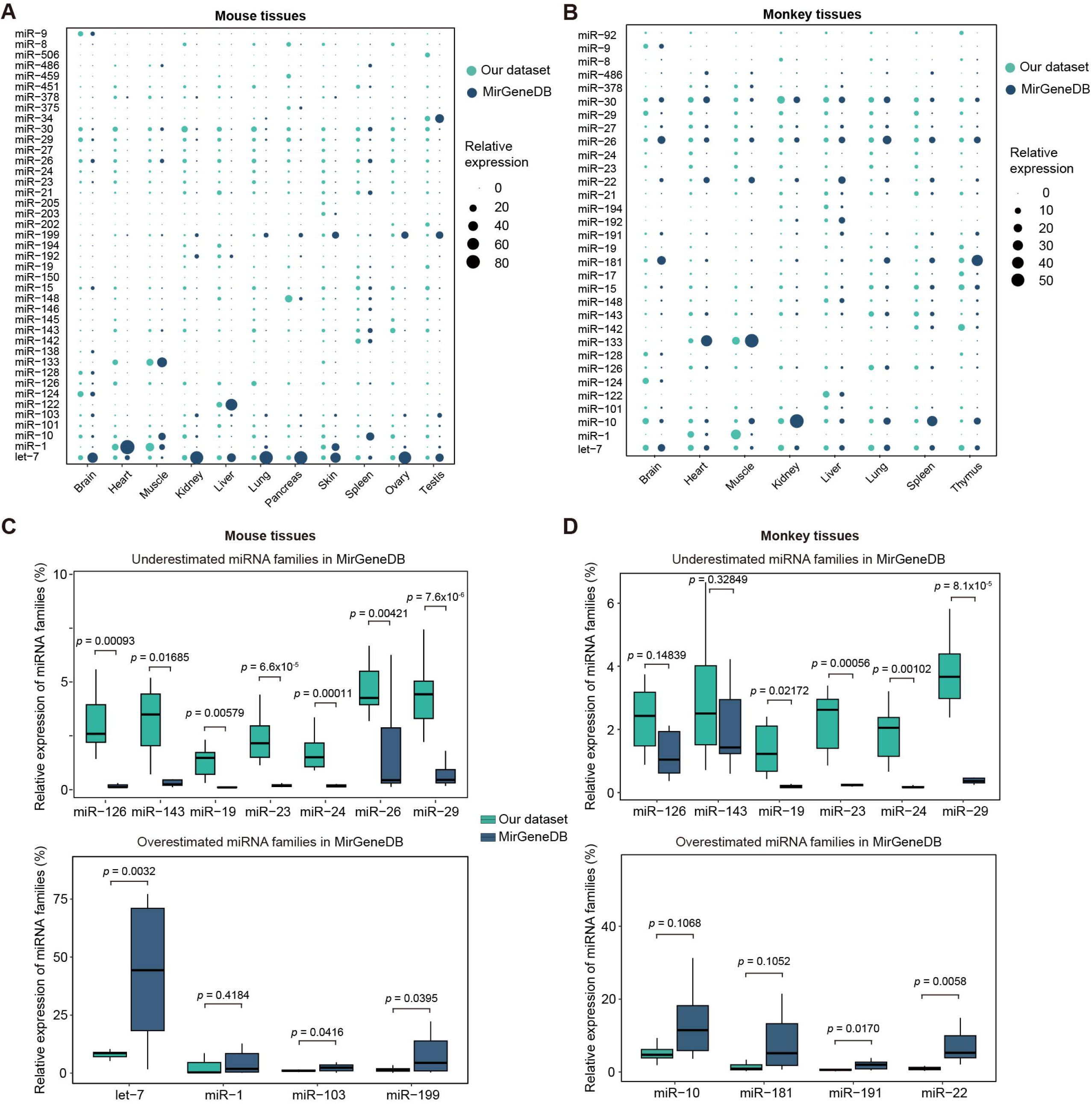
Re-evaluation of miRNA family expression (A-B) Expression patterns of the top 30 miRNA families in mice **(A)** and monkeys **(B)** based on our datasets or the MirGeneDB database. Circle size represents the proportion of miRNA family expression relative to total miRNA families. Biological replicates: n = 2 (for mouse ovary and testis), n = 3 (for mouse-breast, monkey-rectum, and monkey-stomach), or n = 4 (for remaining samples). **(C-D)** Comparison of overestimated or underestimated representative miRNA family expression between our datasets and the MirGeneDB database for mouse **(C)** and monkey tissues **(D)**. Box plots in (C) and (D) depict the median (center line), interquartile range (box), and min-max range (whiskers). Significance was assessed using two-tailed Student’s t-tests.

**Supplementary Figure 5:**
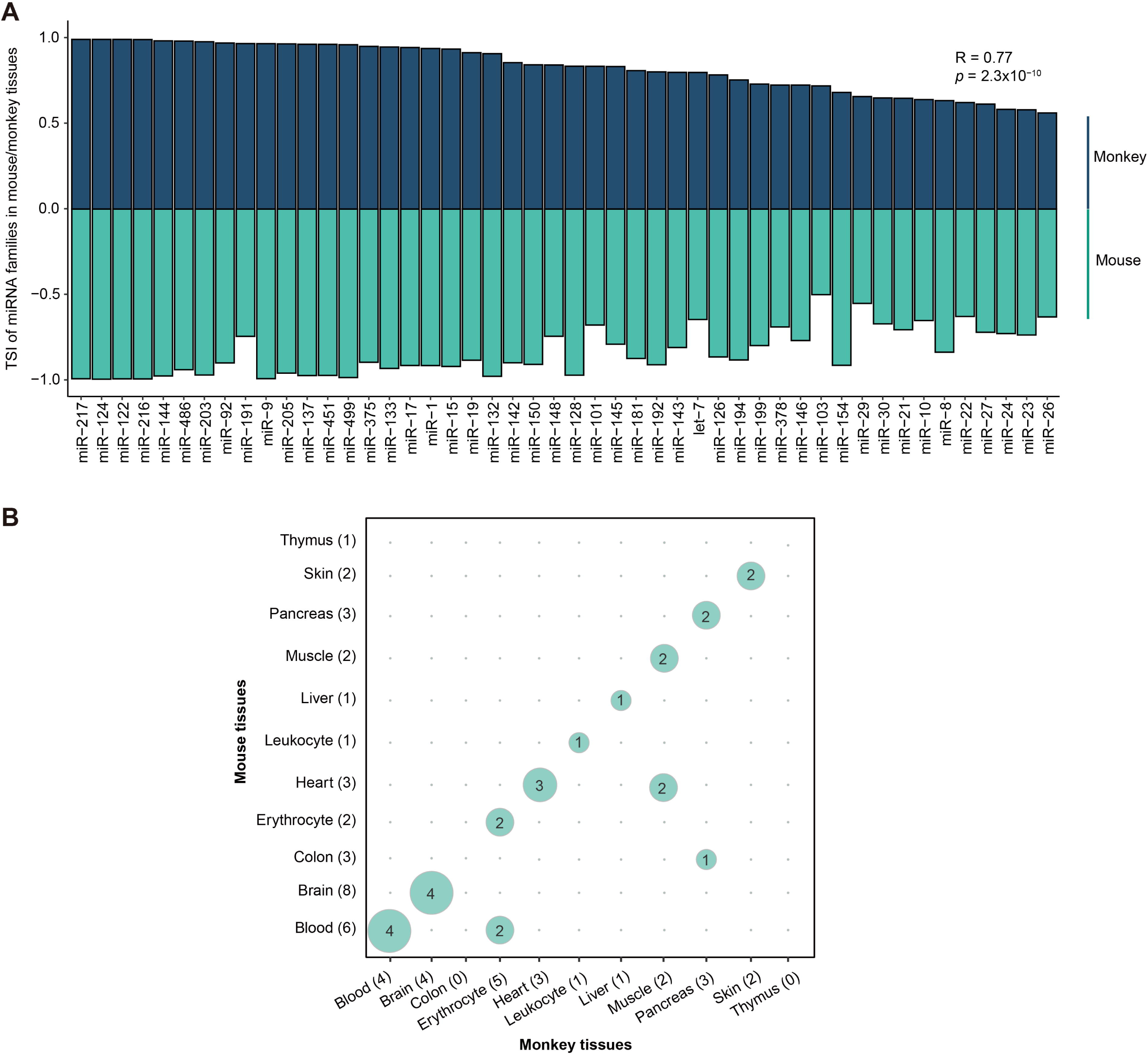
Tissue-specific miRNA families are highly conserved between mice and monkeys. **(A)** TSI distribution of highly expressed miRNA families (the proportion of miRNA family expression relative to total miRNA families >1%) in monkey and mouse tissues. Significance was assessed using two-tailed Student’s t-tests. **(B)** Confusion matrix of miRNA families with a TSI greater than 0.85. Numbers inside the bubbles indicate the number of overlapping tissue-specific miRNA families between monkey and mouse tissues. Numbers in parentheses along the x- and y-axes represent the tissue-specific miRNA families identified in monkeys and mice, respectively.

**Supplementary Figure 6:**
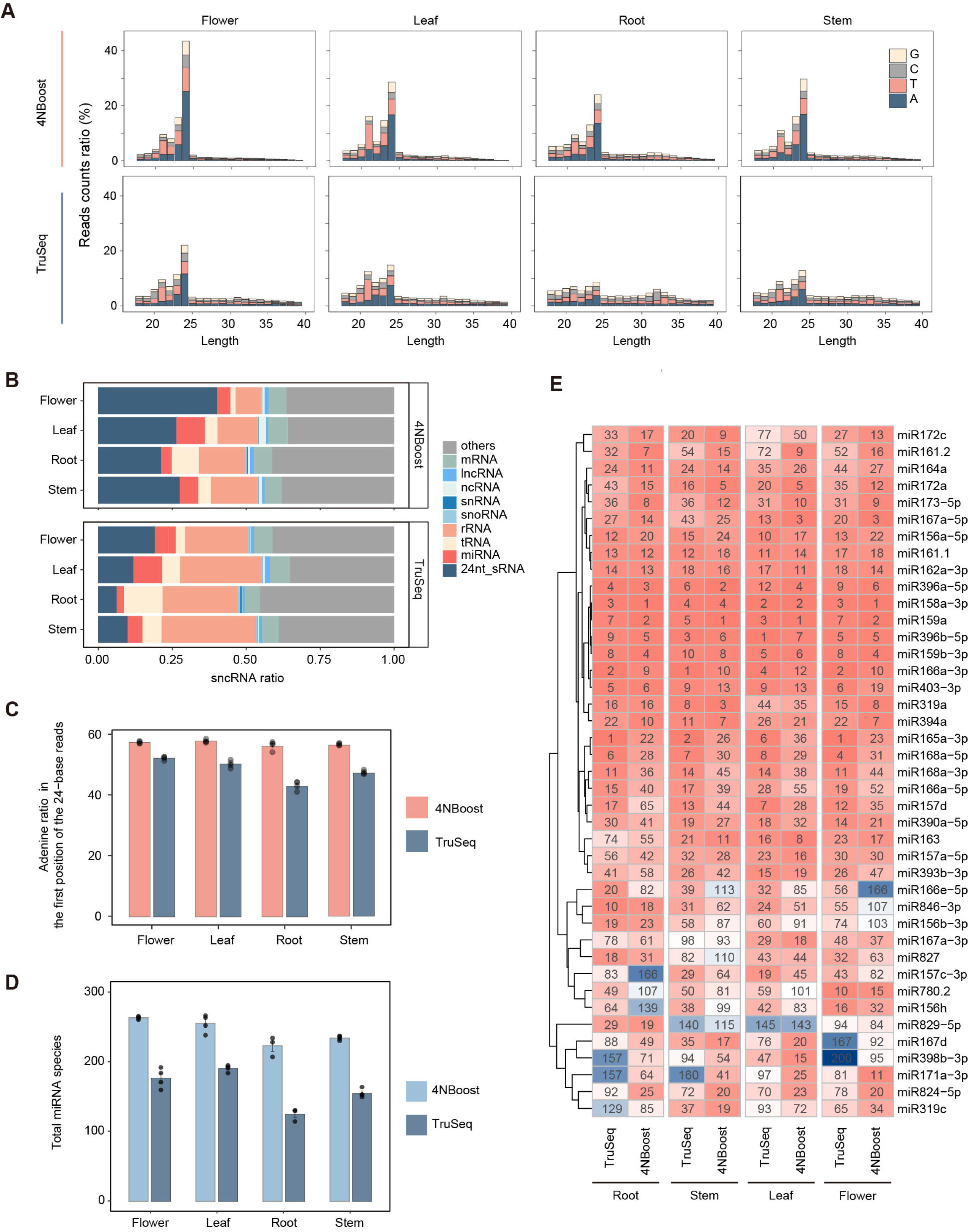
Re-evaluation of sncRNAs abundance in *Arabidopsis thaliana* tissues. **(A)** The distribution of the first nucleotide across sncRNA-seq reads of different lengths. Data are obtained from libraries of four *Arabidopsis thaliana* tissues prepared using either the 4NBoost or TruSeq. Colors indicate the proportion of sncRNA-seq reads beginning with each nucleotide. **(B)** Proportions of various sncRNA types in four tissue libraries prepared using the 4NBoost and TruSeq. **(C)** Relative abundance of 24-nt sncRNAs starting with 5’-adenine in 4NBoost or TruSeq libraries across all four tissues. **(D)** The number of distinct miRNA species detected in each tissue by the two methods. **(E)** Comparison of miRNA expression rankings between libraries prepared using the 4NBoost and TruSeq across all four tissues. All panels are based on the data from three or four biological replicates per tissue. Data in (A), (B), and (E) are presented as mean values. Data in (C) and (D) are presented as mean ± s.e.m.

**Supplementary Table 1.** Representative miRNAs with previously validated strand.

| miRNAs | Cell lines | Previously validated strand preferences |  | Strand preferences in our datasets |
| --- | --- | --- | --- | --- |
|  |  | Methods | Preferences |  |
| Pre-mir-324 | HEK293T | Northern | 5p | 5p |
| Pre-mir-423 | Hela & HEK293T | Northern & RT-qPCR | 3p | 3p |
| Pre-mir-223 | Jurkat | RT-qPCR | 3p | 3p |
| Pre-mir-133a | Hela<sup>1</sup> | Northern | 3p | 3p |

preferences and those identified by our datasets.
1. pre-mir-133a is not naturally expressed in HeLa cells. This cell line was used to transfect constructs that expressed pre-mir-133a to determine its strand preferences.

**Supplementary Table 2.** Methods for cell lysis of each tissue.

| Mouse |  | Crab-eating monkey |  |
| --- | --- | --- | --- |
| Tissue | Method | Tissue | Method |
| lung | magnetic beads | lung | liquid nitrogen |
| breast | liquid nitrogen | breast | liquid nitrogen |
| colon | magnetic beads | colon | magnetic beads |
| rectum | magnetic beads | rectum | magnetic beads |
| kidney | magnetic beads | kidney | magnetic beads |
| brain | magnetic beads | brain | liquid nitrogen |
| skin | liquid nitrogen | skin | liquid nitrogen |
| liver | magnetic beads | liver | magnetic beads |
| stomach | magnetic beads | stomach | magnetic beads |
| spleen | magnetic beads | spleen | magnetic beads |
| pancreas | liquid nitrogen | pancreas | liquid nitrogen |
| heart | magnetic beads | heart | magnetic beads |
| thymus | magnetic beads | thymus | magnetic beads |
| muscle | magnetic beads | muscle | magnetic beads |
| adipose | magnetic beads | adipose | magnetic beads |
| testis | magnetic beads | - | - |
| ovary | magnetic beads | - | - |

**Supplementary Table 3.**
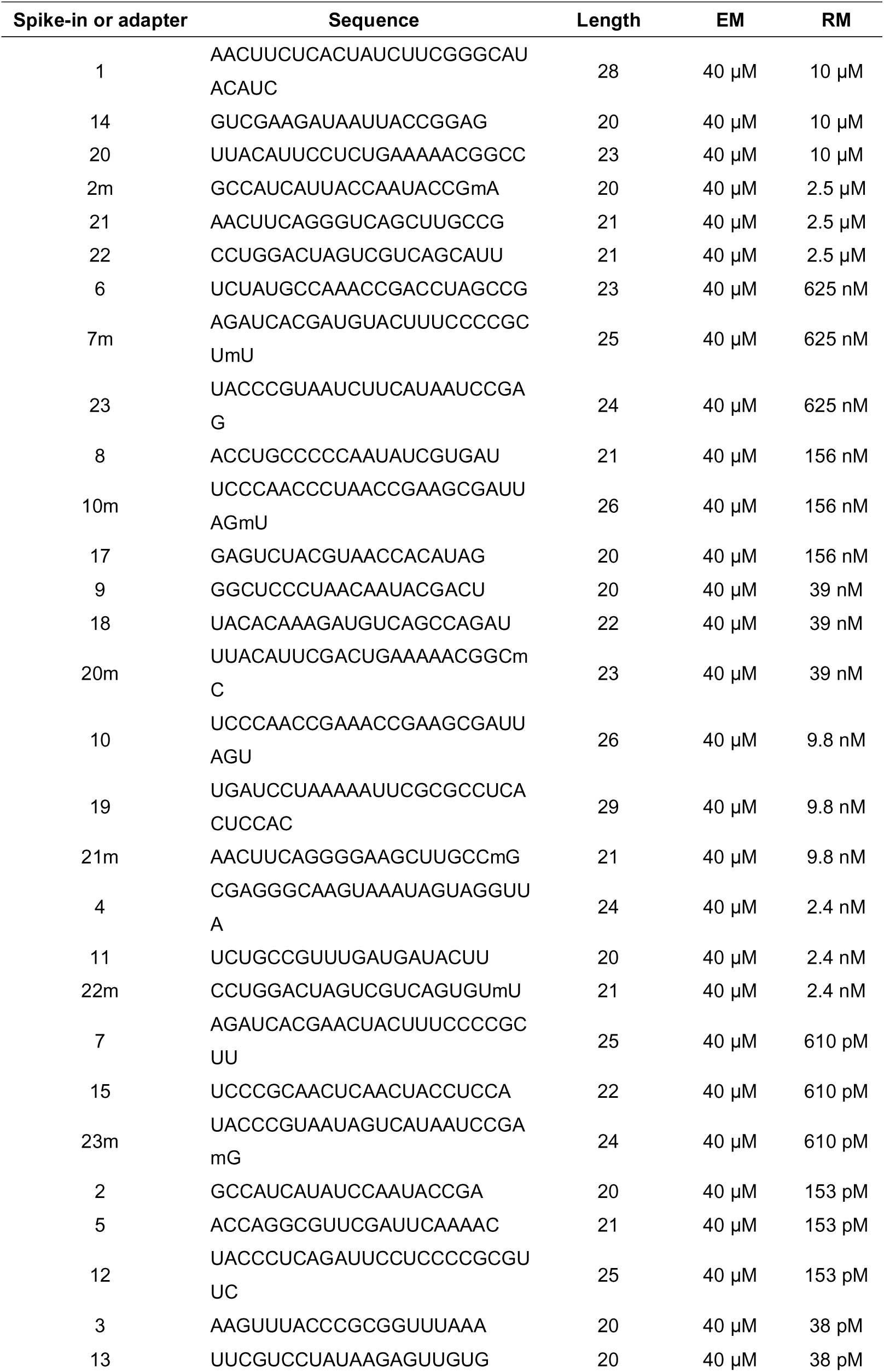
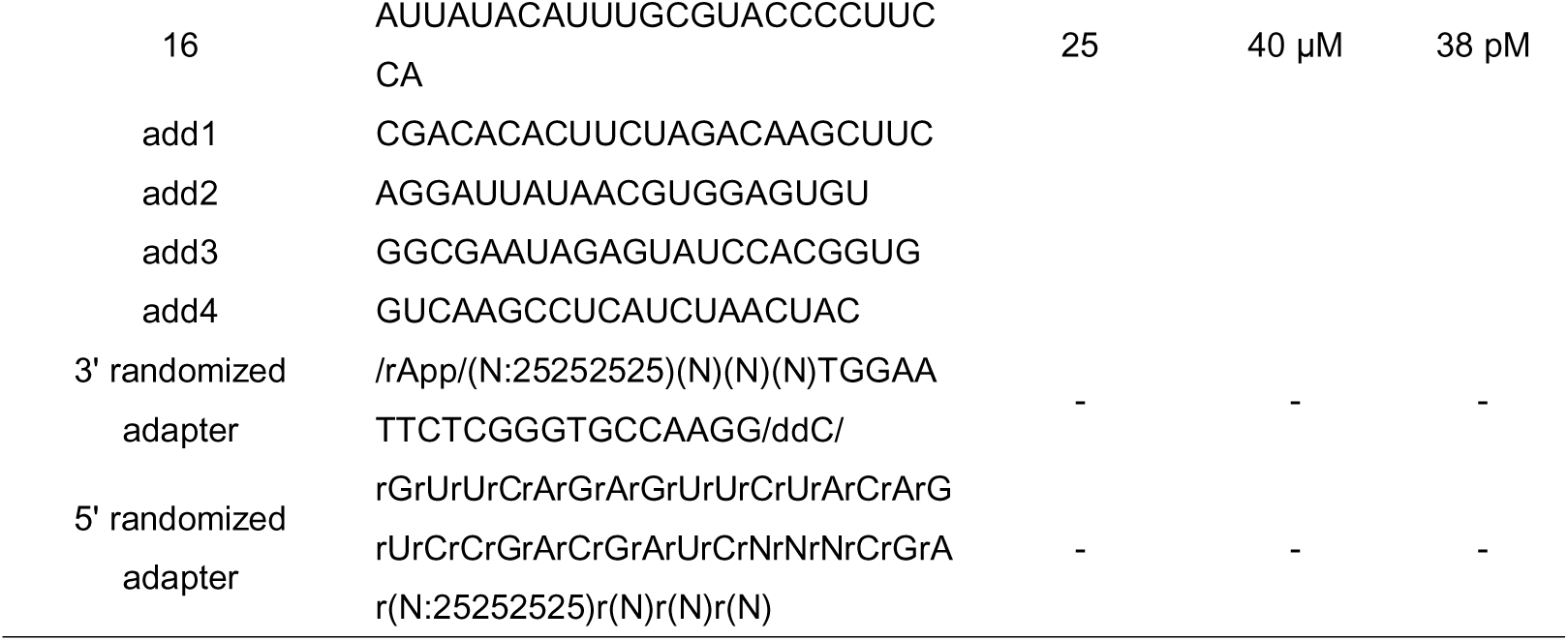
The sequences of adapters, along with the sequences and specific mixing ratios of EM and RM spike-ins.

**Supplementary Table 4.**
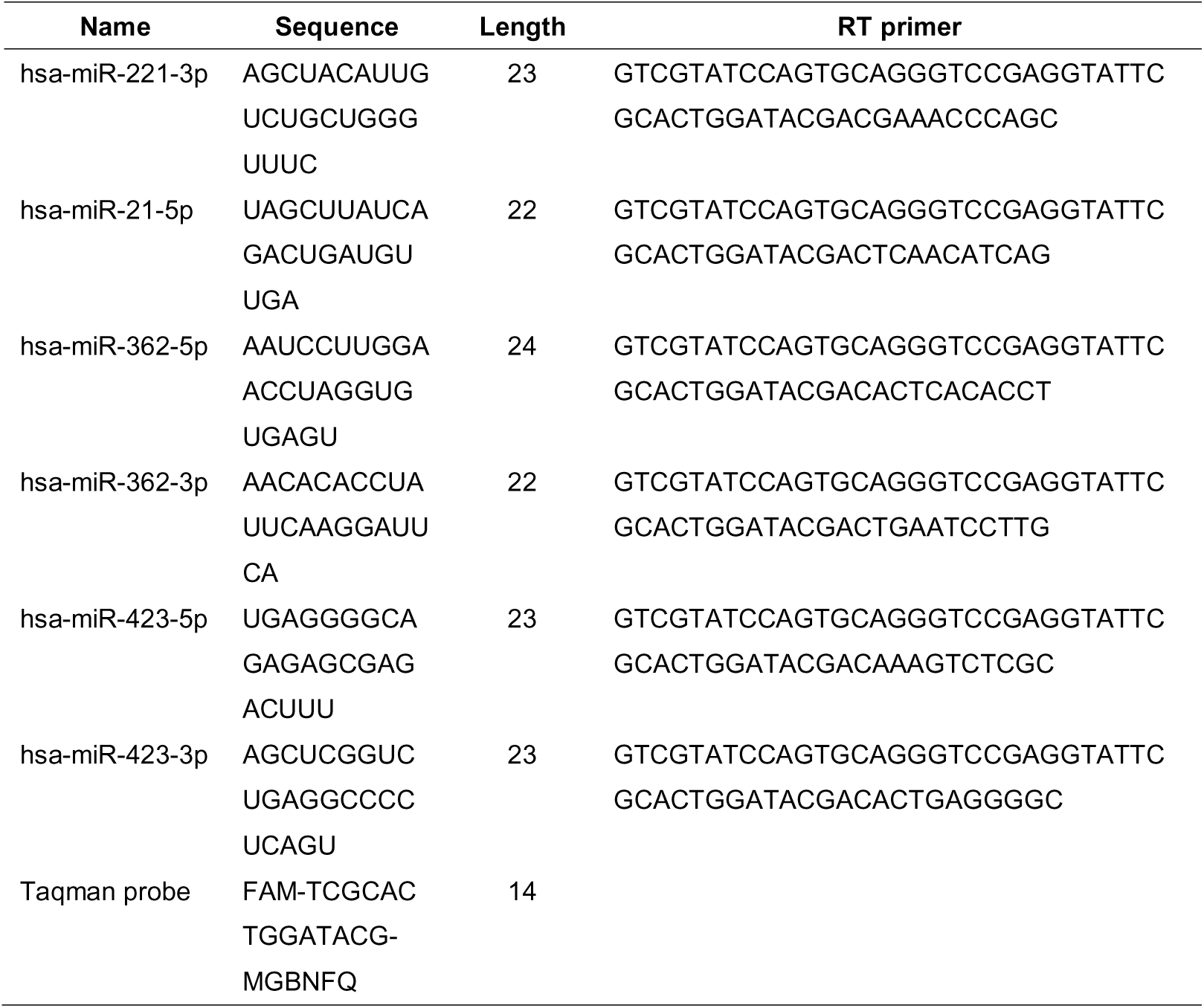
The sequence of RT-qPCR primer.

## Notes

### Competing Interest Statement

The authors have declared no competing interest.

### Summary of Updates

To avoid any nomenclature conflict and ensure clarity, we have renamed our method as 4NBoost in the revised manuscript. Moreover, we rigorously validated the quantification accuracy of 4NBoost using RT-qPCR, and further demonstrated its broader applicability by profiling sncRNAs in Arabidopsis thaliana tissues, thereby highlighting its potential as a valuable tool for sncRNA research in plant biology.

